# Predictability in the evolution of Orthopteran cardenolide insensitivity

**DOI:** 10.1101/542811

**Authors:** Lu Yang, Nitin Ravikanthachari, Ricardo Mariño-Pérez, Riddhi Deshmukh, Mariana Wu, Adam Rosenstein, Krushnamegh Kunte, Hojun Song, Peter Andolfatto

**Author notes:** Correspondence: Lu Yang:, Peter Andolfatto.

## Abstract

The repeated evolutionary specialisation of distantly related insects to cardenolide-containing host plants provides a stunning example of parallel adaptation. Hundreds of herbivorous insect species have independently evolved insensitivity to cardenolides, which are potent inhibitors of the alpha-subunit of Na^+^, K^+^-ATPase (ATPα). Previous studies investigating ATPα-mediated cardenolide insensitivity in five insect orders have revealed remarkably high levels of parallelism in the evolution of this trait, including the frequent occurrence of parallel amino acid substitutions at two sites and recurrent episodes of duplication followed by neo-functionalisation. Here we add data for a sixth insect order, Orthoptera, which includes an ancient group of highly aposematic cardenolide-sequestering grasshoppers in the family Pyrgomorphidae. We find that Orthopterans exhibit largely predictable patterns of evolution of insensitivity established by sampling other insect orders. Taken together the data lend further support to the proposal that negative pleiotropic constraints are a key determinant in the evolution of cardenolide insensitivity in insects. Furthermore, analysis of our expanded taxonomic survey implicates positive selection acting on site 111 of cardenolide-sequestering species with a single-copy of ATPα, and sites 115, 118 and 122 in lineages with neo-functionalised duplicate copies, all of which are sites of frequent parallel amino acid substitution.

## Introduction

Two enduring fundamental questions in modern evolutionary biology are *what factors limit the rate of adaptive evolution?* and *to what extent are adaptive evolutionary paths predictable?* [1] Theoretically, the predictability of adaptation depends on a number of factors that constrain the number of possible evolutionary paths. Among these is the number of potential targets for beneficial mutations [2–4]. However, the extent to which adaptation is constrained by mutation rate is unclear for most traits and some investigators have emphasised important roles for pleiotropy, the phenomenon by which one mutation affects multiple phenotypes, and epistasis, the effect of genetic background on the contribution of a mutation to a given phenotype, among other factors [1,5–7].

Evaluating the relative importance of these factors has been challenging. One approach has been to cobble together examples of adaptations from different traits in different species and contexts in an attempt to come to general conclusions (e.g. [8,9]). However, the heterogeneity inherent in such broad comparisons of different traits in different biological contexts may substantially limit the power to make inferences from such data [10]. An alternative is to examine cases on a trait-by-trait basis in the context of adaptation to common selective pressure [10]. Instances of parallel evolution, the independent evolution of similar features in different lineages, can provide multiple portraits of the evolutionary process and offer insight into the factors that constrain adaptation and the extent to which adaptive evolutionary paths are predictable.

Examples of parallelisms from nature are abundant and occur at different scales from the resemblance of morphological traits to individual nucleotide substitutions that encode regulatory or protein changes [4,11,12]. Such examples will have greater power to make inferences about the factors determining the dynamics of adaptation for a given trait as the number of independent outcomes becomes larger, the more similar the selective pressure and the more is known about the genetic basis of the underlying trait. With these factors in mind, one fertile area for exploration is the repeated evolution of insensitivity of herbivorous insects to toxic secondary plant compounds. Plants are ubiquitously equipped with secondary chemical defences such as alkaloids, cyanogenic glucosides, and terpenoids that contribute to defence against herbivory [13,14]. Despite these defences, herbivorous insects have in many cases repeatedly evolved mechanisms to render them insensitive to toxic compounds [14,15], and even sequester toxins for their own use [16].

A striking example is presented by herbivorous insects that have repeatedly evolved the ability to feed on and, in many cases, sequester cardenolides from Apocynaceae plants, which include milkweed [15]. Cardenolides represent a class of steroidal glycosides (“cardiac glycosides”) that bind to and inhibit the alpha-subunit of Na^+^, K^+^-ATPase (ATPα). This protein is a ubiquitously distributed membrane-bound ion active-transporter present in animals with well-known roles in a variety of physiological processes including neural signal transduction, muscle contraction and osmoregulation [17]. Conservation of the cardenolide-binding domain of ATPα among distantly-related animals, including vertebrates and invertebrates, suggests important physiological roles for the regulation of Na^+^, K^+^-ATPase by endogenously produced cardenolides [17,18]. In fact, cardenolides have been used medicinally for hundreds of years as common treatments for conditions such as congestive heart failure and cardiac arrhythmias [18]. A growing number of studies have implicated the regulation of Na^+^, K^+^-ATPase by putatively endogenous cardenolides in signalling pathways linked to a variety of pathologies including hypertension and cancer [19,20].

Due to its medical importance, the interaction between Na^+^, K^+^-ATPase and cardenolides has been well-studied. The binding of cardenolides arrests Na^+^, K^+^-ATPase in the phosphorylated state, where K^+^ cannot be bound, Na^+^ cannot be released to the extracellular side and ATP is not hydrolysed [21]. Mutagenesis experiments, enzyme-ligand co-crystal structures, and evolutionary analyses have implicated 41 amino acid residues of ATPα, scattered throughout the protein, that either directly interact with cardenolides or affect their binding-affinity indirectly (references listed in **Table S1**). These sites are largely concentrated near the site of cardenolide binding in ATPα, with some exceptions (**Figure S1**). As such, the evolution of cardenolide-insensitivity via the modification of ATPα (i.e. target-site insensitivity) is one of the rare traits for which we have a good *a priori* idea of the beneficial mutation target size for adaptation.

Broadly speaking, strategies employed by specialist herbivores to deal with toxin compounds include destroying and/or excreting the toxins [22,23], inactivating the toxins by chemical modifications [24], restricting the expression of the target protein to specific tissues [25,26], and/or the evolution of target-site insensitivity [24]. The evolution of ATPα insensitivity has so far been inferred in almost all Apocynaceae-specialist species surveyed from five insect orders, including Lepidoptera, Diptera, Coleoptera, Hymenoptera, and Hemiptera [27–33].

Studies of the evolution of ATPα insensitivity in these five insect orders have revealed a remarkable degree of convergence of molecular mechanism at multiple levels. First, despite the identification of 41 residues in the protein that could potentially modulate sensitivity of ATPα to cardenolides (**Table S1**), there is a marked enrichment of substitutions in Apocynaceae-specialists observed at two sites in the protein, Q111 and N122, that flank the H1-H2 extracellular loop [30,33]; the substitutions Q111L, Q111T, Q111V and N122H occur in parallel in multiple lineages. Second, several rounds of duplication of ATPα have occurred in parallel in multiple species from three of the five surveyed insect orders, including Coleoptera, Hemiptera, and Diptera [30,33]. In each case, at least one of the divergent copies harbours a number of cardenolide insensitivity-conferring substitutions [30,33,34]. Third, in most cases of ATPα duplication, the copies have been shown to exhibit parallel evolution of tissue-specific expression patterns, with putatively less cardenolide-sensitive copies predominating in the gut, and putatively more sensitive copies predominating in nervous tissue [30].

Taken together, these data suggest a key role for negative pleiotropy in the evolution of cardenolide-insensitivity in Apocynaceae-specialists: those species with a single copy are largely limited to evolution at just a few of the 41 possible sites, but those with duplicates can explore many other evolutionary paths using one of the differentially expressed duplicate copies [30]. Further support for the idea that negative pleiotropy plays a key role comes from site-directed mutagenesis studies demonstrating trade-offs in Na^+^, K^+^-ATPase function (e.g. efficiency of ATP hydrolysis) associated with some duplicate-specific substitutions observed in milkweed bugs (Hemiptera) [35].

The notion of predictability in evolution is often framed in terms of forecasting future events, for example, predicting the next steps in virus evolution [36]. However, it can also be used in the sense of looking at evolutionary patterns retrospectively and asking if a set of rules deduced from patterns of evolution in one group of organisms for a particular adaptation can reliably predict the genetic architecture of the same trait in another group. With this in mind, we survey a sixth insect order, the Orthoptera, which is phylogenetically positioned outside of the five insect orders that have previously been investigated (**Figure 1A**). Within Orthoptera, we focus on the family Pyrgomorphidae, commonly known as gaudy grasshoppers. This group is a relatively small group of approximately 500 species that include some of the most colourful and showy grasshoppers in the world (**Figure 1B-D**) [37]. Some members of this family that are known to feed on toxic plants, including Apocynaceae, possess aposematic colouration and are able to sequester plant secondary metabolites such as cardenolides and pyrrolizidine alkaloids [38–41]. In addition to the aposematic colouration, some genera (such as *Phymateus, Poekilocerus, Zonocerus*) possess a unique mid-dorsal abdominal gland capable of squirting toxic chemical when disturbed, while others (*Aularches, Dictyophorus, Taphronota*) can produce foam as a result of haemolymph released through pores combined with air [41–44].

**Figure 1.**
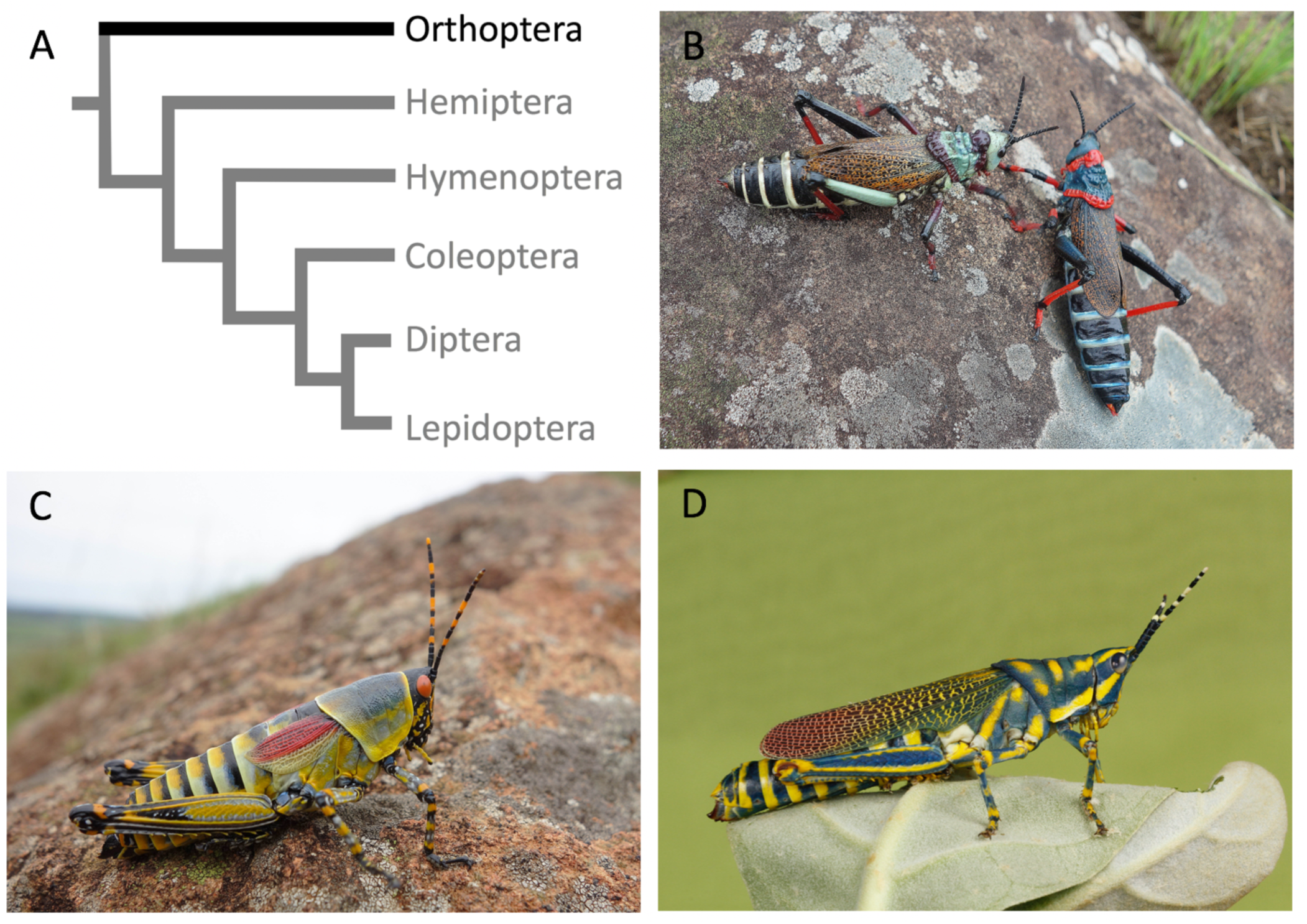
Phylogeny (A) and representative members of the Orthoptera: Pyrgomorphidae (B-D). Orthopteran species shown are B) *Dictyophorus spumans* (South Africa); C) *Zonocerus elegans* (South Africa); D) *Poekilocerus pictus* (India).

Studies of the common milkweed grasshopper *Poekilocerus bufonius* demonstrated that they were substantially less sensitive to the cardenolide ouabain injections (as measured by LD50) than species that do not feed on milkweed plants [45]. In addition, enzyme inhibition assays performed on extracts from *P. bufonius* suggest cardenolide insensitivity of Na^+^, K^+^-ATPase. In addition, observed heterogeneity among tissues in the degree of cardenolide-insensitivity was interpreted as possible evidence for distinct isoforms of the enzyme in this species [45]. While several species within Pyrgomorphidae show plant-mediated chemical defence, most of the species in the family do not possess aposematic colouration or feed on toxic plants [37], which presents an interesting opportunity to investigate the variation in cardenolide-insensitivity of ATPα.

To shed light on the evolution of cardenolide-insensitivity in Pyrgomorphidae, we generated RNA-Seq data for one representative grasshopper species from each of ten genera in the family and reconstructed the ATPα by *de novo* transcriptomic assembly [30]. We find remarkably similar patterns of amino acid substitution to those observed in previous surveys of Apocynaceae-specialist herbivores, including a duplication event followed by neo-functionalisation and tissue-specific differential expression in the genera *Phymateus* and *Poekilocerus*. This expanded dataset, now including data for 52 Apocynaceae-feeding species from six insect orders, further supports the view that adaptation in this system appears to be largely constrained by negative pleiotropic effects associated with otherwise adaptive substitutions. The dataset also affords increased power to detect positive selection acting on specific sites in the protein that are also sites of recurrent parallel amino acid substitution.

## Materials and Methods

### Sequencing and *de novo* transcriptome assembly

For details on sample collection and preparation see **Table S2**. Dissections were carried out in Phosphate-buffered saline solution and stored either in TRIzol (Ambion, Life Technologies) or RNAlater (Ambion Inc.) at −80°C. For all insects, total RNA was extracted using TRIzol (Ambion, Life Technologies) following the manufacturer’s protocol. RNA-seq libraries of *Aularches miliaris* and *Poekilocerus pictus* were prepared with TruSeq Stranded total RNA Library Prep Kit with Ribo-Zero Gold (Illumina) and sequenced on Illumina HiSeq2500 at AgriGenome (Cochin, Kerala, India). The libraries of *Taphronota calliparea* and *Dictyophorus griseus* were prepared with NEBNext Ultra RNA library Preparation Kit (NEB) and sequenced on Illumina HiSeq4000 (Genewiz, South Plainfield, NJ, USA). The libraries of *Chrotogonus hemipterus*, *Atractomorpha acutipennis*, *Zonocerus elegans*, *Phymateus leprosus*, *Ochrophlebia cafra*, and *Sphenarium purpurascens* were prepared with TruSeq RNA Library Prep Kit v2 (Illumina) and sequenced either on Illumina HiSeq4000 (Genewiz, South Plainfield, NJ, USA) or HiSeq2500 (Genomics Core Facility, Princeton, NJ, USA). Reads were trimmed for adapters, quality and length (Phred quality ≥20 and ≥30 contiguous bases) using TQSfastq.py (http://code.google.com/p/ngopt/source/browse/trunk/SSPACE/tools/TQSfastq.py). All transcriptomes were *de novo* assembled with Trinity 2.2.0 [46]. ATPα of *Locusta migratoria* (GenBank: KF813097.1) was used to query the assembled transcripts using BLAST (blast-2.26). Reconstructions of ATPα for each species were used iteratively as query sequences to BLAST against each other using either tblastx or blastn to recover all ATPα copies.

We have also included previously unpublished full-length ATPα sequences for a number of other Apocynaceae-specialists including *Daphnis nerii* (Oleander Hawk-moth, Lepidoptera), *Empyreuma pugione* (Spotted Oleander Caterpillar Moth, Lepidoptera), *Euploea core* (the Common Crow, Lepidoptera), *Danaus chrysippus* (Plain Tiger, Lepidoptera), *Liriomyza asclepiadis* (Milkweed Leaf-Miner Fly, Diptera). The methods to reconstruct these sequences is identical to those used above following Zhen *et al*. [30]. Particular attention is paid to 41 sites implicated in cardenolide-insensitivity (**Table S1**) established based on site-directed mutagenesis and protein-ligand co-crystal structure studies.

### Discovery and confirmation of duplicates

Given previous studies revealing duplications of ATPα associated with Apocynaceae-specialisation [30], we evaluated evidence for duplication in our Orthopteran species data. Two ancient lineages of ATPα (ATPα1 and ATPα2) precede the diversification of multiple insect orders and form a distinct clade from the duplications of ATP1A found in vertebrates [30]. However, the expression level of ATPα2 is low in insects surveyed to date and its function remains largely unknown. In *Drosophila melanogaster*, expression of ATPα2 is limited to larval imaginal discs and adult male testes and accessory glands; RNAi and P-element knock-outs of the gene are homozygous viable but male-sterile (http://www.flybase.org/reports/FBgn0267363). Furthermore, we failed to recover an ortholog of ATPα2 from the *Locusta migratoria* genome assembly [47], and several other Orthopteran assemblies [48,49], suggesting that the ATPα1/α2 duplication may have arisen after the split of Orthopterans from the other insects we have surveyed (**Figure 1A**). We thus decided to focus on ATPα1, which shows clear orthology across taxa and a strong correlation between evolution at known cardenolide-sensitivity sites in the protein and specialisation on cardenolide-containing Apocynaceae plants [30].

The discovery of duplicated copies of ATPα1 in *Poekilocerus pictus* and *Phymateus leprosus* was verified by cloning and sequencing. Total RNA was extracted as described above, and reverse-transcribed to single-strand cDNA using SuperScript^®^ III Reverse Transcriptase (Thermo Fisher Scientific). ATPα1 was PCR-amplified using Phusion High-Fidelity DNA Polymerase (Thermo Fisher Scientific) using forward primer: 5’-ACATGGCGGCAAGAAGAAAG-3’ and reverse primer: 5’-AGTAGGGGAAGGCACAGAAC-3’. The PCR product was cleaned using QIAquick PCR Purification Kit (Qiagen), 3’A-tailed using Taq polymerase (NEB) and cloned into TOPO TA vector (Invitrogen) following the manufacturer’s instructions. Ampicillin-resistant colonies were picked and screened by colony-PCR for the presence of inserts on a 1% agarose gel. Libraries of plasmids were constructed using Tn5 transposase [50] annealed with Tn5ME-A, 5’-TCGTCGGCAGCGTCAGATGTGTATAAGAGACAG-3’ and Tn5ME-B, 5’-GTCTCGTGGGCTCGGAGATGTGTATAAGAGACAG-3’ and indexed with customised Illumina i7, i5. Paired-end 150 nt reads were collected for the pooled library on an Illumina MiSeq Nano (Genomics Core Facility, Princeton University). *De novo* transcriptome assembly was performed with Velvet/Oases [51,52] using a random sub-sample of 10,000 reads for each indexed plasmid.

### Differential expression analysis

Head, muscle, and foregut tissues of three male and three female *Poekilocerus pictus* were dissected and sequenced in two batches (see sample collection and sequencing). Adapter and quality trimmed reads were mapped back to our *de novo* assembly of the transcriptome as a reference (including ATPα1A and ATPα1B) using bwa mem [53] with default criteria, processed with SAMtools 0.1.18 [54] and mapped reads were counted with HTSeq 0.6.1 [55]. We used the inverted beta-binomial (ibb) test [56] to determine the significance of difference of expression level between tissues. The method uses a negative binomial distribution in a generalised linear model framework for paired-sample testing. We applied a standard Bonferroni correction to account for multiple tests. Paired-sample count data were normalised by either total number of mapped reads or the sum of reads mapping to ATPα1A and ATPα1B (**Table S3**).

### Re-analysis of Lygaeid ATPα1 evolution and expression

Using the recently completed *Oncopeltus fasciatus* genome [57], we detected a fourth copy of ATPα1 (ATPα1D) in these Lygaeid bugs that was missed by previous studies. Lygaeid ATPα1D is the least-derived copy at sites implicated in cardenolide-sensitivity and is expressed in *O. fasciatus* heads, yet has the lowest expression of the four copies (**Figure 3A**). We also confirmed that, like copies A-C, ATPα1D is also shared with *Lygaeus kalmii* (a sister-genus species) but we could only partially reconstruct it from our *de novo* transcriptome assembly due to its low expression. Using RNA-seq data for *O. fasciatus*, we confirmed the finding of differential expression of duplicates documented by Zhen *et al.* [30] **(Figure S3A)**. All four copies of ATPα1 in the Lygaeid *O. fasciatus* are more highly expressed in the head than the gut. The putatively most sensitive copy (ATPα1D), despite having the lowest expression level of the four copies, exhibits the greatest degree of up-regulation in the head relative to other copies (**Figure S3B**).

### Evolutionary analyses

The ages of the duplicates were calculated from dS (the per site rates of substitution at synonymous sites) estimated using PAML4.8 codeml [58], with prior trees based on established cladistic relationships (Mariño-Pérez *et al*, unpublished, [59])and calibrated with divergence times obtained from www.timetree.org (*Locusta* and Pyrgomorphidae: 117.4 Mya; *Napomyza* and *Phytomyza*: 39 Mya). The divergence times of the Large Milkweed Bug (*Oncopeltus fasciatus*), Milkweed Stem Weevil (*Rhyssomatus lineaticollis*), and Dogbane Beetle (*Chrysochus auratus*) are taken from [30] where similar methods were used to date duplicates. In the Lygaeid bugs (*O. fasciatus* and *L. kalmii)*, a phylogenetic analysis strongly suggests a duplication order ((ABC),D), ((AB),C) and most recently (A,B).

To obtain distributions of lineage-specific evolutionary rates, ATPα1 lineages were grouped into four categories. ATPα1 of all non-specialist species were denoted as "Outgroup". ATPα1 of Apocynaceae-specialists with a single copy of ATPα1 are denoted "Single". For specialists with multiple copies of ATPα1, copies that are up-regulated in the gut relative to the head are assumed to be relatively cardenolide-insensitive copies and marked as Dup^I^ [30].

Likewise, those up-regulated in the head relative to the gut are assumed to be relatively sensitive copies and were grouped as Dup^S^. We chose this criterion rather than the number of insensitivity-conferring substitutions because using the latter makes the designation of copies with intermediate numbers of substitutions ambiguous. In Lygaeid bugs (*O. fasciatus* and *L. kalmii)*, this implies that copies A and B are treated as relatively insensitive copies (Dup^I^), whereas C and D are treated as relatively sensitive (Dup^S^) (**Figure S3**). The dN/dS ratios (omega) for ATPα1 along a lineage were estimated within each insect order using PAML codeml under free ratio model. Parameters were set as follows: seqtype = 1, model = 1, NSsites = 0, clock = 0, CodonFreq = 2, fix_kappa = 0, kappa = 2.0, fix_omega = 0, omega = 0.02. Differences in the distribution of branch-specific estimates of omega between each group were tested with Dunn’s test of multiple comparisons using rank sums as implement in R (dunn.test).

We also evaluated evidence for positive selection acting on individual sites of ATPα1 using PAML codeml. To do this, we defined and investigated several models (**Figure 7, Table S4**). Model 1: positive selection on all Apocynaceae-specialist lineages; Model 2: positive selection on all Apocynaceae-specialist lineages with single copies of ATPα1 (including ancestral lineages prior to duplication); Model 3: positive selection on Dup^S^ and Dup^I^ lineages of Apocynaceae-specialists; Model 4: positive selection on Dup^I^ lineages of Apocynaceae-specialists; Model 5: positive selection on all outgroup lineages. Tests were carried out using the modified branch-site model A implemented in codeml [60–62] with parameters set as follows: seqtype = 1, model = 2, NSsites = 0, clock = 0, CodonFreq = 2, fix_kappa = 0, kappa = 2.0, fix_omega = 0, omega = 0.02. The ancestral sequences of duplicated ATPα1 were reconstructed with the function RateAncestor. Unrooted trees were used, and branch labels were added manually for each model. The method assigns each codon a <u>B</u>ayes <u>E</u>mpirical <u>B</u>ayes (BEB) posterior probability that the codon belongs to a site class with omega > 1 (i.e. indicating positive selection). A BEB posterior probability of >0.95 was considered evidence for positive selection.

## Results

### Survey of ATPα1 of 15 Orthopteran genera

Using an RNA-seq-based gene discovery method, we reconstructed the complete coding sequences of the alpha subunit of Na^+^, K^+^-ATPase (ATPα1) of grasshoppers from species representing ten genera in the family Pyrgomorphidae, as well as five outgroup species within, and one outside, the order Orthoptera (**Figure 2**). Our broad survey of Apocynaceae-feeding Pyrgomorphidae revealed few amino acid substitutions among the 41 sites implicated in cardenolide-sensitivity (**Table S1**). The two most broadly distributed substitutions, Q111L and A119S, correlate only weakly with Apocynaceae-feeding, aposematism, and the presence of abdominal defensive glands in the group. The glaring exception is the lineage leading to the genera *Poekilocerus* and *Phymateus*, both containing multiple species, which appear to share a duplication of ATPα1. Both species surveyed retain an ancestral version of the protein (ATPα1B) while having a highly-derived copy (ATPα1A). The diverged ATPα1A copies of *Poekilocerus* and *Phymateus* share many amino acid substitutions relative to the ancestral copy, several at sites implicated in cardenolide-sensitivity. Phylogenetic analysis clearly indicates that the duplication of ATPα1 and functional divergence of ATPα1A predates the diversification of these clades into separate genera and species and we estimate the age of the duplication to be ∼36 million years old.

**Figure 2.**
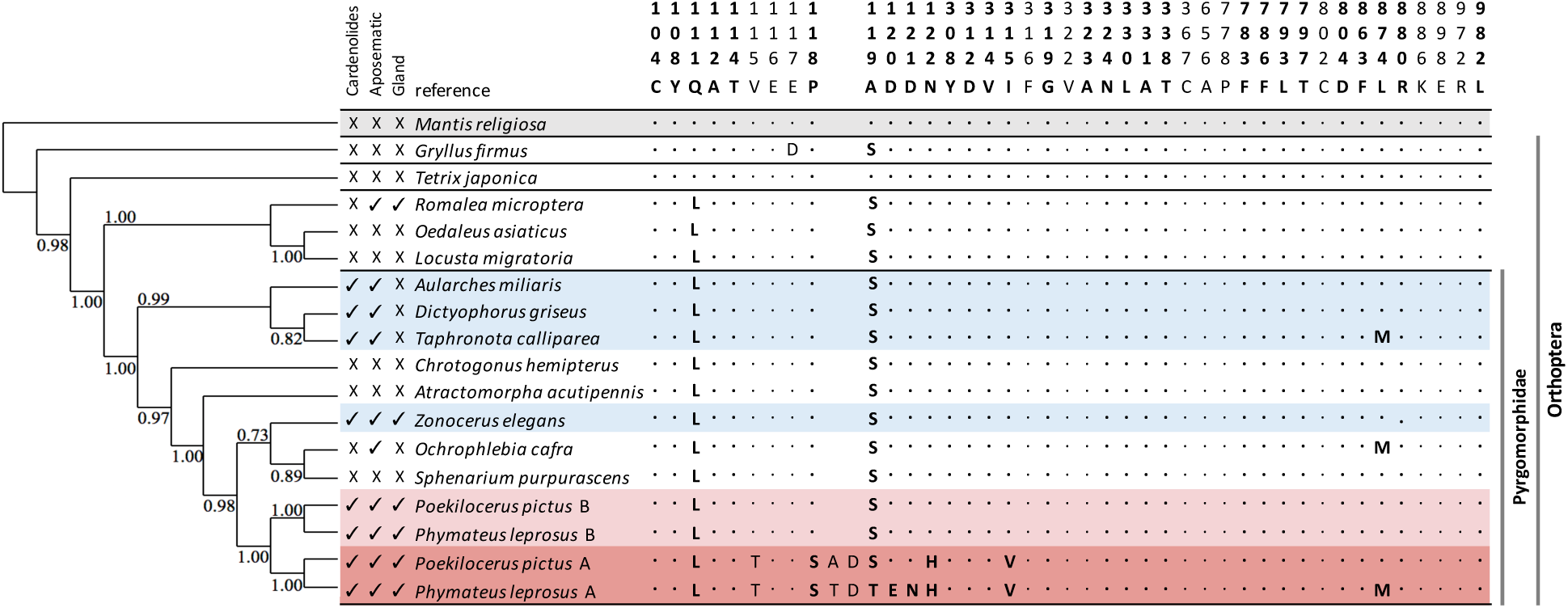
Amino acid substitutions at sites implicated in cardenolide-sensitivity for ATPα1 of Orthopterans in the family Pyrgomorphidae and outgroups. Each family is separated by black lines. The coloured rows correspond to putatively cardenolide-adapted species that either possess only one copy (light blue) or two copies (light/dark red) of ATPα1. The non-Orthopteran outgroup (*Mantis religiosa*, order Mantodea) is shaded in grey. The numbering of sites is based on sheep ATP1A1 (*Ovis aries*) (Genbank: NC019458.2). Bold columns correspond to the sites for which site-directed mutagenesis studies suggest a role in cardenolide-sensitivity, while the rest were identified by structural prediction. Dots indicate identity with the reference, which represents the consensus sequence among non-specialist Arthropods. Letters represent amino acid substitutions relative to the reference. The cladogram was constructed using a maximum likelihood method implemented in SeaView based on protein-coding nucleotide sequences of ATPα1 with bootstrap values shown. “Cardenolide” refers to species feeding on Apocynaceae. “Aposematic” refers to the presence of warning colouration patterns. “Gland” refers to the presence of a specialised abdominal defensive gland that secrete toxic chemicals.

### Patterns of amino acid substitution in ATPα of Orthopterans

Cross-referencing the pattern in Orthopterans with other Apocynaceae-specialists surveyed in other insect orders reveals a high level of parallel amino acid substitution (**Figure 3, Figure S2**). Conspicuous among these are two parallel substitutions Q111L and A119S, which appear to pre-date the diversification of the Pyrgomorphidae. Cell transfection experiments have shown when substitutions at position 111 are introduced to a sensitive background of the *D. melanogaster* protein, the survival rate of HeLa cells increases 3 to 8-fold [31]. Interestingly, A119S is observed in almost every Apocynaceae-specialist species that has been surveyed to date (**Figure S2**), including the *Drosophila subobscura* subgroup where resistant forms of ATPα1 have been documented segregating as polymorphisms within *D. subobscura* [63].

**Figure 3.**
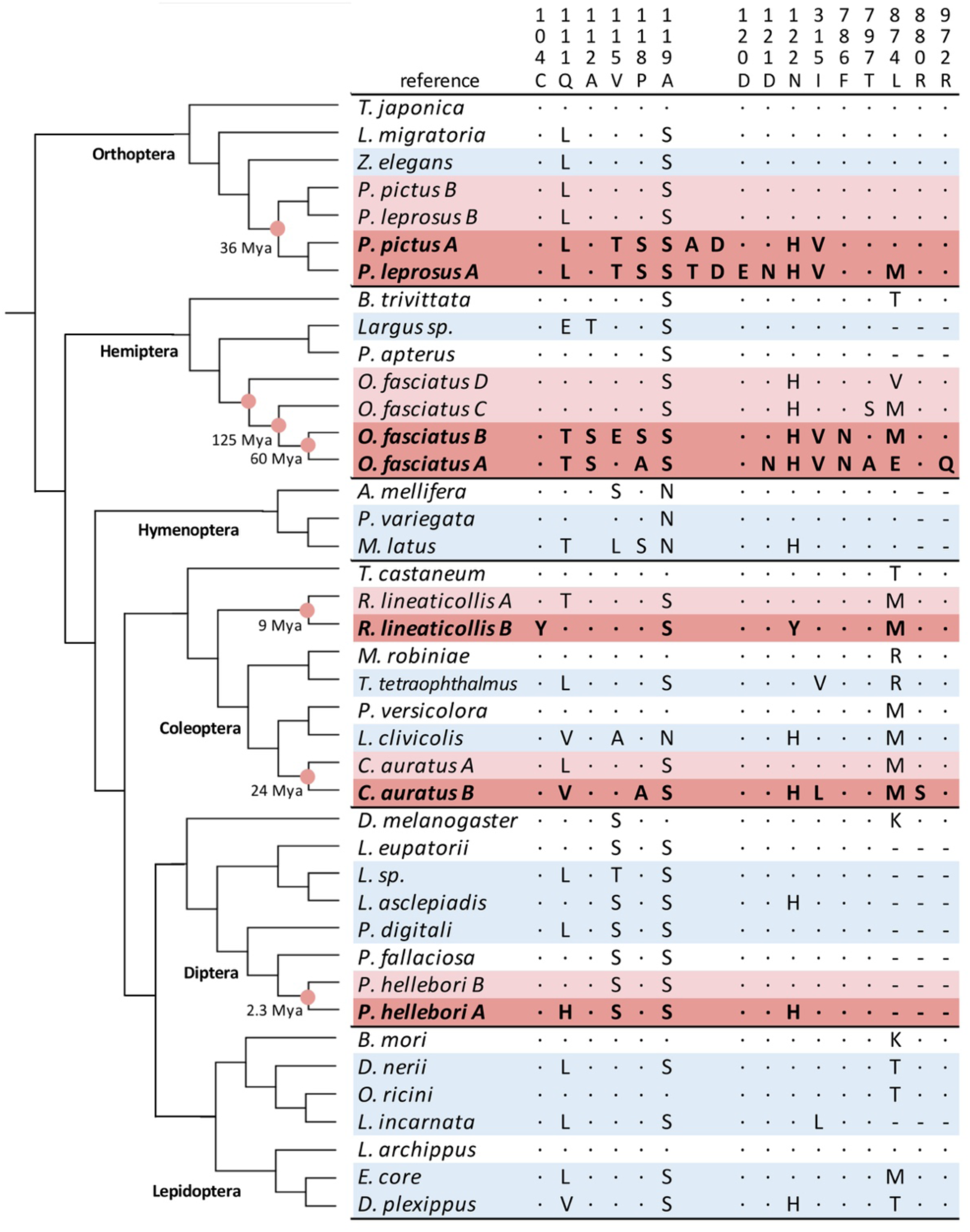
Patterns of amino acid substitution and duplication spanning six insect orders. Shown are sites implicated in cardenolide binding (**Table S1)** at which substitutions have occurred. The colour scheme is consistent with **Figure 2**. Red circles indicate duplication events. The cladogram represents the relationships of insect orders and species phylogenies (see Methods).

Exceptions include aphids (*Aphis nerii* and *Acrythosiphon pisum*), the milkweed leaf beetle (*Labidomera clivicolis*) and several Hymenopteran species that harbour the similar substitution, A119N. Q111L and A119S are not associated with cardenolide-feeding or Apocynaceae-specialisation in the Orthoptera and are possessed by a number of non-aposematic species not known to feed on cardenolide-containing plants (e.g. *L. migratoria*, *C. hemipterus*, *A. acutipennis*). The substitutions Q111L, A119S, and A119N also occur sporadically among a number of other insects not known to feed on Apocynaceae (**Figure S2**).

Considering the cardenolide-insensitive ATPα1A copy-specific substitutions of *Poekilocerus* and *Phymateus*, N122H stands out as a substitution observed in parallel in many other Apocynaceae-specialists including at least one member of each of the six insect orders surveyed. The N122H substitution in isolation has been shown to increase *Drosophila* ATPα1 insensitivity to the cardenolide ouabain by 250-fold [64] and increase survival of HeLa cells challenged with ouabain [31]. N122H has also been reported to interact synergistically with substitutions at Q111, though Q111L was not tested [31]. The Orthopteran ATPα1A copy-specific substitutions V115T, P118S, D121N, I315V and L874M also occur in at least one other insect order. Of these, D121N is known to decrease cardenolide-sensitivity by ∼100-fold [65].

### Unique substitutions associated with duplication of ATPα1 and differential expression of neo-functionalised copies

Using data from three insect orders, Zhen *et al.* [30] found a significant enrichment of unique substitutions at sites implicated in cardenolide-sensitivity in Apocynaceae-specialist lineages with duplicated copies of ATPα1 compared to those that retain a single copy. They also documented convergent patterns of differential gene expression of independently derived duplicates found in specialists. Specifically, they noted that duplicates inferred to be the most sensitive to cardenolides, based on the number of substitutions at sites implicated in cardenolide-insensitivity, consistently exhibited up-regulation in the head relative to the gut. Zhen *et al*. [30] argued that this might be expected since the gut is the site of first-processing of cardenolides and sensitive forms of ATPα1 in nervous tissue are likely protected to some extent by the blood-brain barrier provided by the glial sheath surrounding neurons [30,66]. Zhen *et al.* [30] interpreted this pattern as being consistent with a key role for pleiotropy in the evolution of cardenolide-insensitivity. Specifically, they proposed that there might be trade-offs in enzyme performance associated with unique substitutions that are ameliorated by differential expression of neo-functionalised duplicate copies. Here, we re-evaluate this claim in the context of our expanded dataset, which now includes Orthoptera and previously published data for five other insect orders.

Despite the remarkable parallelism at the levels of gene duplication and amino acid substitutions, it is notable that “unique” substitutions (i.e. those unique to one lineage) at sites implicated in cardenolide-sensitivity in Orthoptera are restricted to duplicated copies, as observed in other Apocynaceae-specialists with duplications. Notably, ATPα1A of *Poekilocerus* and *Phymateus* share a unique two amino acid insertion between residues 119 and 120 of the ancestral protein (**Figure 2**). Insertions or deletions have not been observed so far among the other five insect orders surveyed to date (**Figure 3, Figure S2**). Furthermore, *Phymateus leprosus* harbours an additional amino acid substitution (D120E) that also appears to be unique among Apocynaceae-specialists. Notably, no unique substitutions were observed among the other 12 Orthopteran species surveyed that appear to retain a single copy of ATPα1.

We also find that the functionally diverged duplicates of ATPα1 in the Orthopteran *Poekilocerus pictus* are differentially expressed in a similar manner to those found in Lygaeid bugs (see Methods) and other Apocynaceae-specialists [30]. Specifically, the putatively less cardenolide-sensitive copy (ATPα1A) is up-regulated relative to the more sensitive copy (ATPα1B) in the gut compared to muscle and head (**Figure 4**). The cardenolide-sensitive copy ATPα1B accounts for 18.7% of total ATPα1 expression in the brain but only 2.2% in the foregut, which is the primary location of cardenolide processing. This finding is consistent with previous studies of the related *Poekilocerus* species, *P. bufonius*, that noted Na^+^, K^+^-ATPase activity of the brain was more sensitive to cardenolide-inhibition than that of the gut [45]. Since the duplication of ATPα1 appears to predate the diversification of *Poekilocerus*, it is highly likely that the tissue-specific ouabain-sensitivity can be explained by the presence of the same duplication we have documented here.

**Figure 4.**
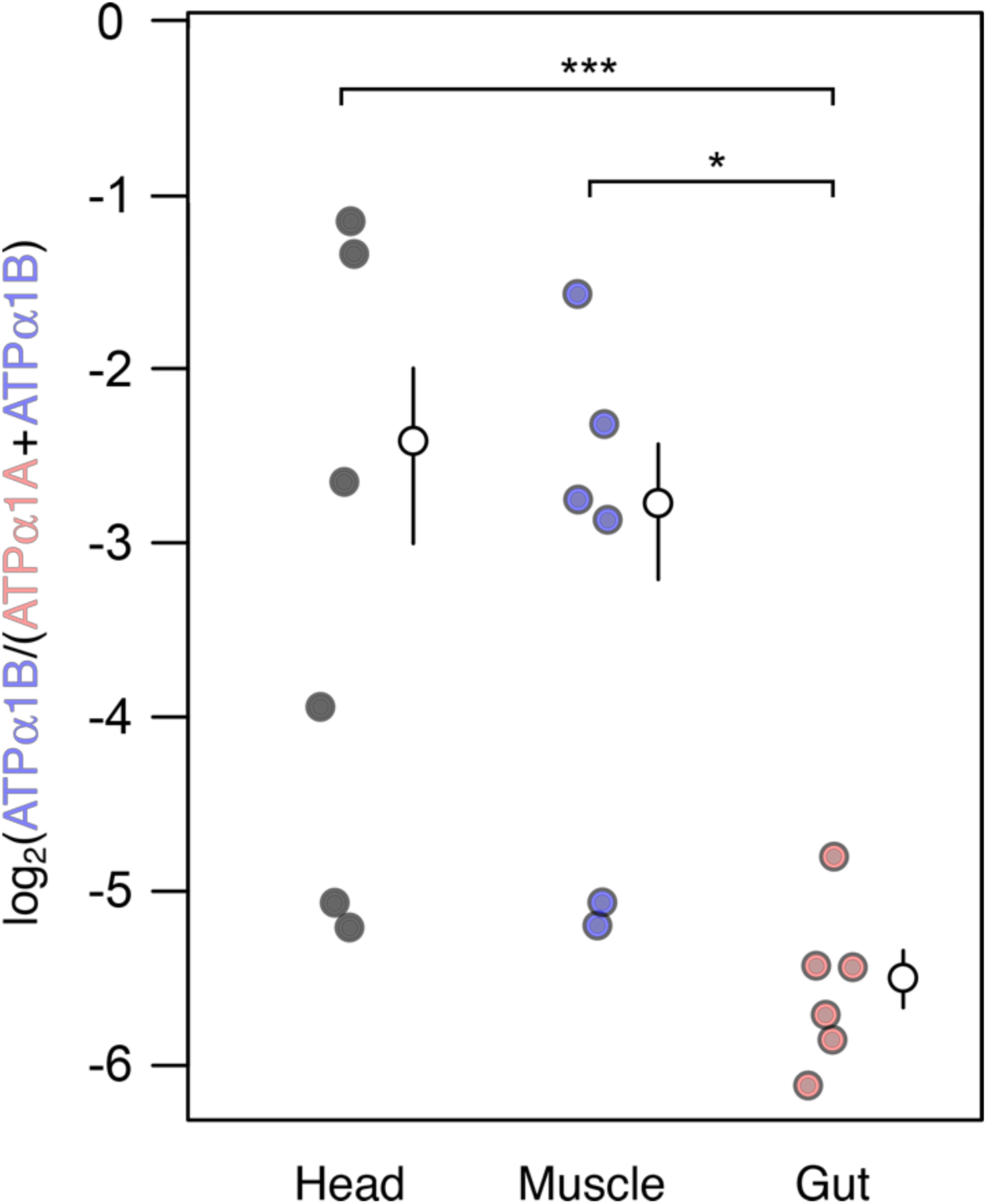
RNAseq-based estimates of ATPα1 expression by tissue in the duplication-bearing Orthopteran *Poekilocerus pictus*. As observed in other Apocynaceae-specialists (Zhen *et al*. [29]), the relatively more cardenolide-sensitive ATPα1B is significantly less expressed in the gut than in head (corrected *P* = 8e-4) and muscle (corrected *P* = 9e-4) in *P. pictus* (**Table S3**). The mean proportion of ATPα1B of total ATPα1 is indicated with open circles with two standard errors as whiskers. P-values were estimated using the “inverted beta-binomial” test for paired sample count data (see Methods).

Having established tissue-specific expression of duplicate copies in Lygaeids (see Methods) and Orthopterans, we then re-visited the pattern of unique versus parallel substitution with respect to duplication status and differential expression of ATPα1 in the full dataset now spanning six insect orders (**Figure 5**). Consistent with Zhen *et al*. [30], we find a marked enrichment of unique substitutions at sites implicated in cardenolide-sensitivity in Apocynaceae-specialists with duplicated copies of ATPα1 (Fisher’s exact test P=0.0022). Examining the pattern in more detail, it is apparent that unique substitutions appear to be unequally distributed among copies. In each case of duplication, we can distinguish between less-sensitive and more-sensitive copies based on the number of derived amino acid-substitutions that have been implicated in cardenolide-sensitivity. We find a significant enrichment of unique substitutions in the dataset (13/14) occur on putatively less-sensitive copies of ATPα1 that are substantially more expressed in the gut than more-sensitive copies (in the 4/5 cases where expression patterns have been investigated).

**Figure 5.**
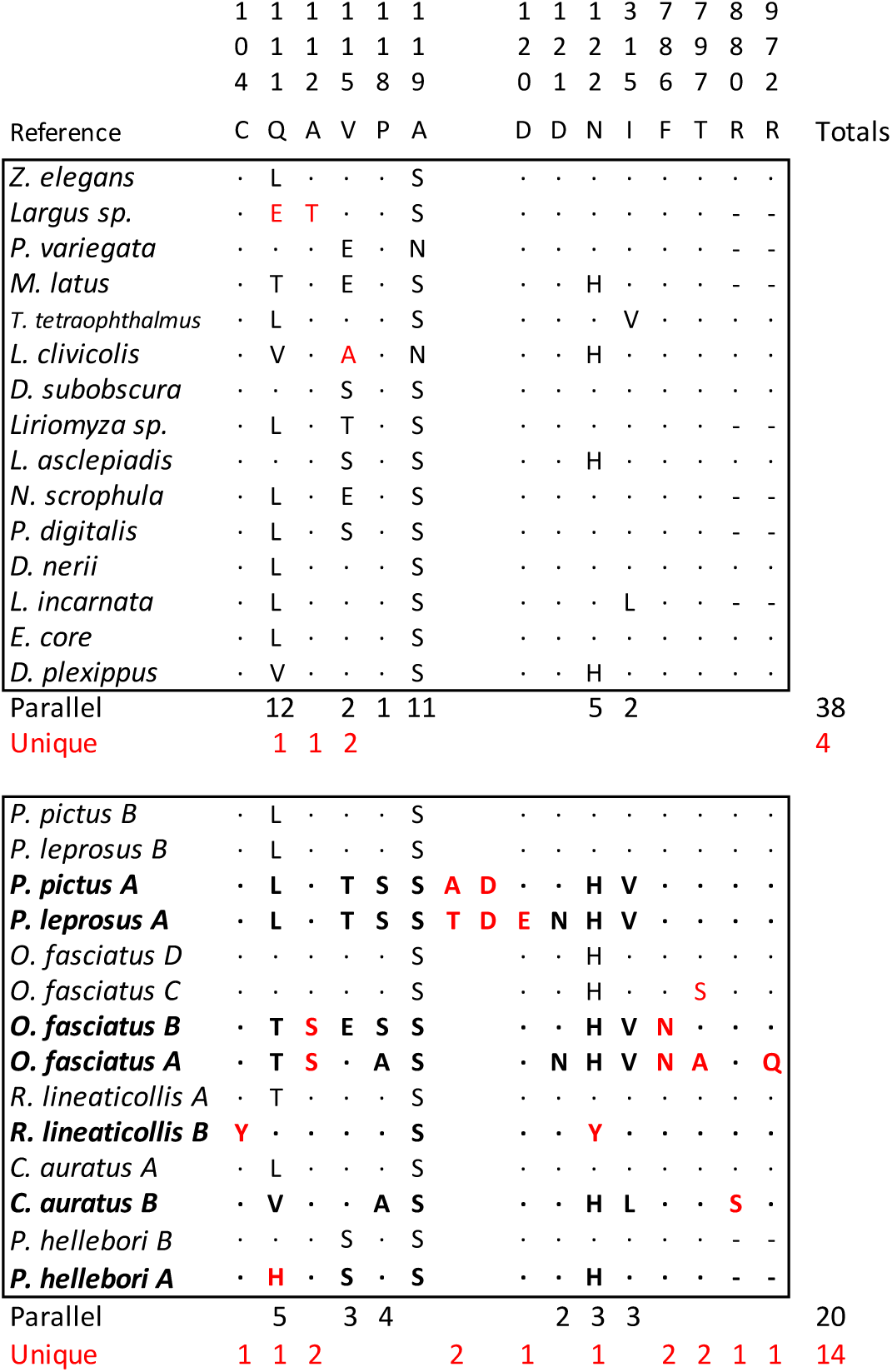
The pattern of parallel versus unique substitutions at sites implicated in cardenolide-sensitivity with respect to duplication status of ATPα1. Unique substitutions are indicated in red. Relatively less cardenolide-sensitive duplicate copies of ATPα1 are highlighted in bold. Site 874 was excluded from this analysis due to uncertainty in the reconstruction of ancestral states.

### Relaxed constraints on ATPα1 duplicates and positive selection for insensitivity

It is clear from the above analyses that the evolution of cardenolide-insensitivity in some taxa is facilitated by duplication and differential expression of ATPα1, which is expected to relax constraints at sites known to confer insensitivity but are associated with negative pleiotropic effects. We further carried out a phylogenetic analysis to ask 1) whether this relaxation in constraint extends beyond sites directly implicated in cardenolide insensitivity, and 2) whether there is evidence for positive selection associated with Apocynaceae-specialisation at these and other sites in the protein. To examine patterns of constraint in more detail, we grouped both external and internal branches of the ATPα1 phylogeny into four categories: Outgroup; Apocynaceae-specialist lineages with a single ATPα1 copy (Single), and Apocynaceae-specialist lineages harbouring duplications that are inferred to be either relatively sensitive (Dup^S^), or relatively insensitive (Dup^I^) to cardenolides (see Methods). Examining the distributions of omega (dN/dS) estimates among these classes reveals that putatively less sensitive copies of ATPα1 (Dup^I^) evolve ∼5-fold faster than their more sensitive counterparts (**Figure 6**). We find that this pattern persists if we exclude the sites directly implicated in cardenolide-sensitivity (**Figure S4**), implying that relaxation of constraint on derived copies extends beyond this class of sites in the protein.

**Figure 6.**
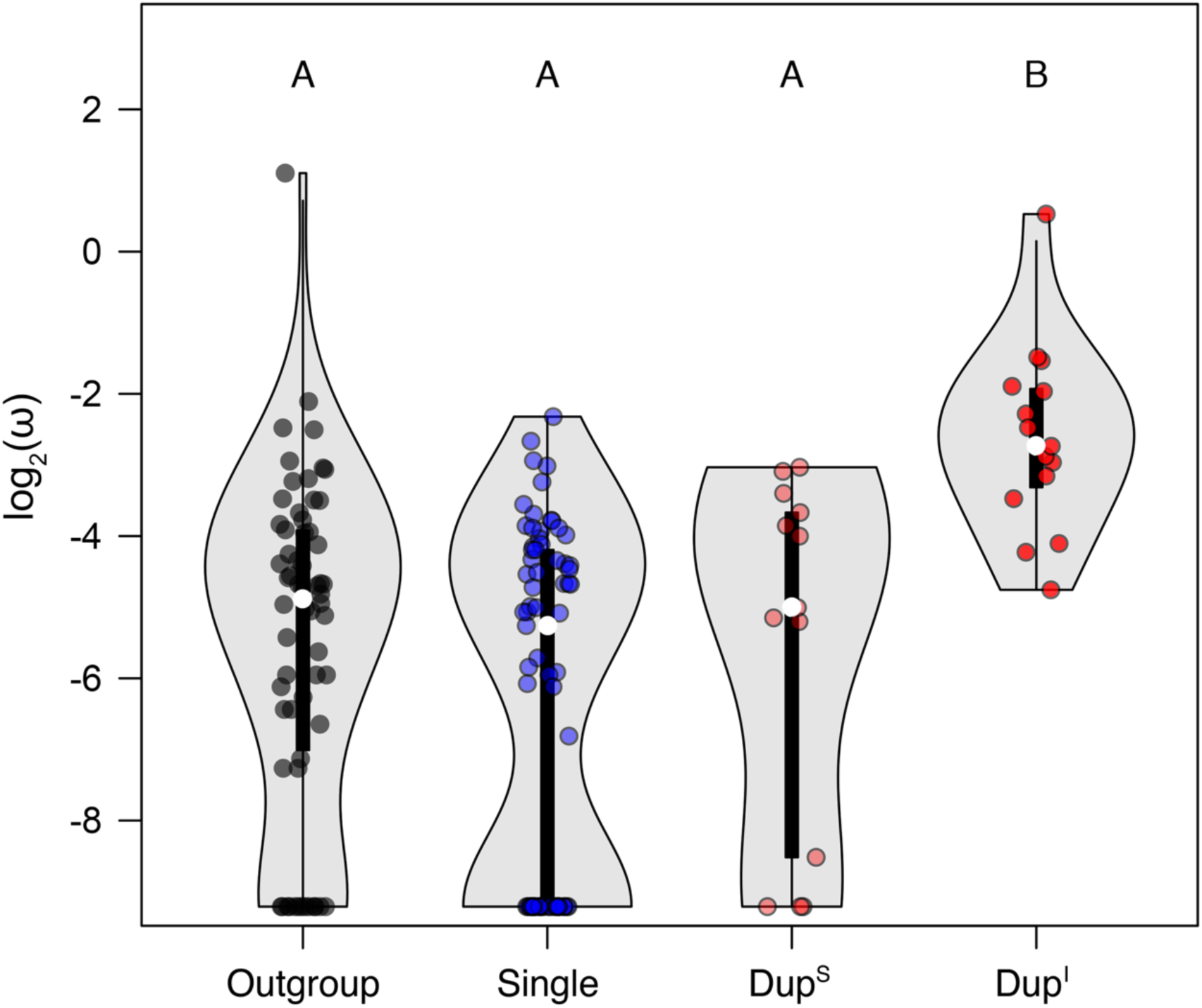
The distributions of omega (dN/dS) estimates for ATPα1 of non-specialists (Outgroup), Apocynaceae-specialists with a single copy of ATPα1 (Single), and specialists with duplicated ATPα1, where the relatively cardenolide sensitive and insensitive copies are noted as Dup^S^ and Dup^I^, respectively. The median omega values are indicated with an open circle and bars represent 50% quantiles. There is a significant difference between the omega distributions for Dup^I^ and those of all three other groups (Letters A and B indicate significantly different categories; Dup^I^ vs Outgroup P=2e-5, Dup^I^ vs Single P=3e-7, Dup^I^ vs Dup^S^ P=3e-3). P-values were estimated using Dunn’s test of multiple comparisons using rank sums as implement in R (dunn.test) and adjusted using the Benjamini-Hochberg method.

We next asked whether relaxed constraint in Apocynaceae-feeding lineages, or on insensitive duplicate ATPα1 lineages, is sufficient to account for the data or whether there is evidence for positive selection associated with insensitivity-conferring amino acid substitutions. We conducted a site-specific scan for positively selected substitutions using the improved branch site model implemented in PAML (see Methods). Site 111, a site directly implicated in cardenolide insensitivity and the target of frequent parallel amino acid substitution across insect orders, is identified as positively selected in this analysis under models assuming either positive selection in Apocynaceae-specialists, or only on Apocynaceae-specialist lineages bearing a single copy of ATPα1 (BEB posterior probability ≈ 1.0 under both models, **Figure 7**, **Table S4**). Sites 115, 118 and 122, also directly implicated in cardenolide-sensitivity, are identified as positively selected under models of positive selection on the Dup^I^ lineages of ATPα1 (BEB posterior probabilities of 0.956, 0.998 and 0.993, respectively, **Figure 7**). Interestingly, evidence for positive selection also emerges at several sites in the protein not previously implicated in cardenolide-sensitivity. Seven sites show evidence for positive selection in lineages bearing duplicate copies of ATPα1 including a cluster of three sites (560, 563, and 566) that are located far from the cardenolide-binding domain (**Table S4, Figure S1**).

**Figure 7.**
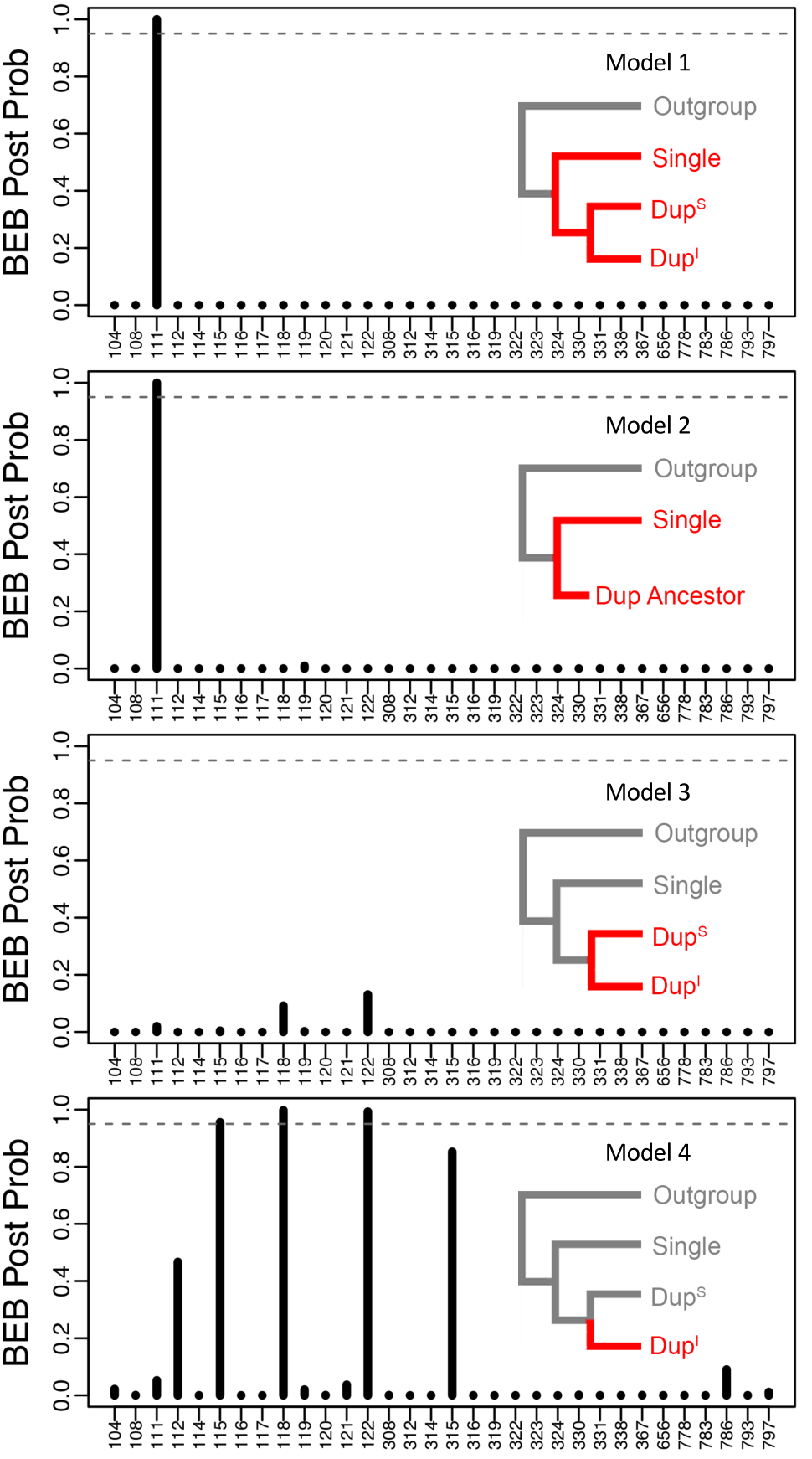
Positively selected sites of ATPα1 among 41 sites implicated in cardenolide-sensitivity (**Table S1**). Sites 802-982 were excluded from the analysis due to missing data. Models 1-4 were tested to identify sites experiencing lineage-specific positive selection. The schematic diagram of each model is shown to the right in each panel. Foreground lineages where positive selection took place are coloured in red and corresponding background lineages are grey. BEB posterior probability >0.95 (grey dashed line) is considered to be strong evidence for positive selection. See **Table S5** for the list of positively selected sites across the whole ATPα1 protein under each model.

## Discussion

Predictability in evolutionary biology not only refers to being able to forecast future evolutionary events but is also a statement about the ability to predict the genetic basis of adaptations outside a taxonomic group in which the rules governing the genetic basis of a particular adaptation were deduced. Previous surveys of ATPα1 in the context of insect adaptation to cardenolides established that, despite a reasonably large target size for evolving target-site insensitivity (i.e. 41 sites implicated in cardenolide-sensitivity, **Table S1**), a small proportion of these sites (and sites 111 and 122 in particular) are disproportionately used in the evolution of insensitivity to cardenolides [30]. The exception to this pattern is found in insects that have duplicated ATPα1 and differentially allocate a functionally-diverged copy to the gut, while retaining an ancestral copy that is more highly expressed in the head.

To test the generality of these patterns as predictors, we surveyed aposematic grasshoppers that sequester cardenolides belonging to Orthoptera, which is the first Polyneopteran order to be examined for cardenolide insensitivity (**Figure 1**). Of the 10 species of Pyrgomorphidae included in the analysis, six species have been reported to feed on Apocynaceae and exhibit aposematic colouration and chemical defence (**Figure 2**). The remaining four species feed on non-toxic plants and show cryptic colouration. While we initially expected all Apocynaceae-feeding species to show substitutions at sites implicated in cardenolide-insensitivity, we found that most Orthopteran species harbour only two substitutions (Q111L and A119S), that are not correlated with consumption and/or sequestration of cardenolide-containing plants in this group. The specific substitution Q111L appears to only weakly confer insensitivity to cardenolides based on previous functional studies. Furthermore, A119S does not interact with cardenolides directly or have an effect on cardenolide-affinity, although it does contribute to more rapid association/dissociation kinetics of the cardenolide ouabain [67]. Given this and their broader phylogenetic distribution (**Figure 3**, **Figure S2**), it is possible that these ancient and recurrent substitutions represent exaptations [68] that facilitate the evolution of more resistant forms of ATPα1 in some insect lineages [29].

Nevertheless, two of the six species (*Poekilocerus* and *Phymateus*) do share a duplication (ATPα1A) that exhibits multiple amino-acid substitutions known to confer insensitivity to cardenolides. Why amino acid substitutions with large effects on cardenolide sensitivity or insensitive duplicates of ATPα1 are not found in the other four Apocynaceae-feeding species deserves an explanation. Of the six Apocynaceae-feeding species included in this study, *Phymateus*, *Poekilocerus*, and *Zonocerus* have the mid-dorsal abdominal glands used for chemical defence [41] and they form a monophyletic group within the phylogeny of Pyrgomorphidae [37]. So far, cardenolides have been reported in the defence secretion from only *Phymateus* and *Poekilocerus* [41], and their common names, milkweed grasshoppers, suggest a strong association of these insects with the host plant. Although *Zonocerus* do feed on cardenolide-containing plants, their main defence chemical is pyrrolizidine alkaloid [69–71], rather than cardenolides. *Zonocerus variegatus* only excretes cardenolides when the toxins are included in its diet [72]. The other three Apocynaceae-feeding species (*Aularches*, *Dictyophorus*, and *Taphronota*) do not have specialised abdominal glands, but produce toxic foams through various pores on the thorax and abdomen [41]. It is unknown whether they use cardenolides as their main chemical defence. The lack of cardenolide insensitivity of Na^+^, K^+^-ATPase in *Zonocerus*, *Aularches*, *Dictyophorus*, and *Taphronota* seems to suggest that they rely on other unknown mechanisms to cope with the toxicity of Apocynaceae. It is possible that while they are capable of consuming Apocynaceae, they might prefer a mixture of other toxic plants containing different kinds of secondary compounds to confer toxicity. Though the chemical ecology of these insects is under-studied, we can postulate that, at least in Pyrgomorphidae, only those species that are intimately associated with Apocynaceae have evolved the cardenolide insensitivity of Na^+^, K^+^-ATPase.

Considering the full dataset now including data from six insect orders, our study confirms previous findings [30] suggesting that, when ATPα1 has been duplicated, unique substitutions at known functionally important sites are significantly enriched specifically among insensitive copies (**Figure 5**). This pattern of relaxation of constraint associated with one copy is also apparent at sites outside of those known to be functionally important sites (**Figure S4**). Both patterns are a strong indication of neo-functionalisation rather than merely sub-functionalisation of ATPα1. This may be expected given the age of the duplication event. Sub-functionalisation is expected to occur rapidly after duplication while neo-functionalisation takes place over longer timescales because it generally requires more mutations [73]. Most of the duplications of ATPα1 in Apocynaceae-specialist insects detected so far are indeed ancient and trans-specific (**Figure 3**).

Despite the expected signature of strong purifying selection on ATPα1 (**Figure 7**, **Table S4**), we have also detected the signature of positive selection on sites implicated in cardenolide-insensitivity and exhibiting frequent parallel substitution. Zou and Zhang [74] have pointed out that the observation of parallel substitution *per se* is not sufficient to infer adaptive significance since this pattern may be expected under a neutral model due to among-site differences in physicochemical constraints. Some evidence against this argument in the case of the evolution of ATPα1 target-site insensitivity is provided by the fact that Apocynaceae-specialist species are highly enriched for substitutions at sites implicated in cardenolide-sensitivity compared to non-specialist outgroups [30; this study]. Our finding of positive selection at some of these sites establishes a more direct link between adaptive protein evolution and recurrent parallel substitutions at sites implicated in cardenolide-sensitivity.

Interestingly, we have also detected signatures of positive selection at sites not previously implicated in cardenolide-sensitivity (**Table S4, Figure S1**). Some of these sites (e.g. 301, 560, 563, 566 and 667) are located far from the cardenolide-binding domain of the enzyme and sites 560, 563 and 566 are known to have roles in binding ATP. While this might seem to preclude direct roles in cardenolide-insensitivity, the existence of sites exhibiting allosteric effects on sensitivity is not unprecedented (e.g. 367 and 656, **Table S1**, [75]). This being said, the detection of positive selection at these sites may reflect selection pressures that are either only indirectly related or even unrelated to the evolution of cardenolide-insensitivity. Future functional experiments could be aimed at understanding the effects of these substitutions on cardenolide-insensitivity and overall enzyme performance.

Duplications of ATPα1 feature prominently in the evolution of cardenolide-insensitivity in herbivorous insects. Given this feature, it will be interesting to compare patterns of recurrent parallel amino acid substitution in insects with vertebrates. Bufonid toads are among the few animals able to produce Na^+^, K^+^-ATPase-inhibiting compounds called “bufadienolides” that closely resemble cardenolides and act in the same way [76]. As a result, predators of bufonid toads represented by a wide variety of vertebrates are under pressure to evolve insensitivity to these compounds [77,78]. In contrast to insects, vertebrates retain at least three copies of ATPα, (ATP1A1, ATP1A2, and ATP1A3), that are differentially expressed among tissues [79].

Previous studies have so far investigated the H1-H2 extracellular loop of ATP1A1 in bufonid toads and predatory frogs [77] and ATP1A3 of a number of predatory squamates [78,80]. It will be of considerable interest to further compare patterns of molecular evolution of cardenolide-sequestering insects to their bufonid predator analogs in the context of complete reconstructions of all three proteins.

## Acknowledgements

We thank Patrick Reilly for help with *de novo* assemblies of transcriptomes and reconstruction of ATPα1. We thank Shurong Hou for constructive comments on protein structure inferences. We thank Piotr Naskrecki (Gorongosa Restoration Project, Mozambique) for help in sample collection. This research was funded by NIH grant R01-GM115523 to P.A. Specimen collection and wet lab work in India was funded by a research grant from NCBS to K.K. Specimen collection in South Africa was funded by NSF grant DEB-1655097 to H.S.

## Data accessibility

All raw RNA-seq sequence data generated for this study have been deposited in the National Center for Biotechnology Information Short Read Archive, www.ncbi.nlm.nih.gov/sra (BioProject PRJNA509040). Sequences of ATPα1 used in our analysis have been submitted to Genbank under Accession numbers MK294065-81.

## Author contributions

L.Y. and P.A. designed the study; N.R., R.M-P, R.D., and K.K. provided the samples; L.Y. performed the experimental work and analysed the data; M.W. contributed to Figure S3; A. R. contributed to Figure S2; L.Y. and P.A. wrote the manuscript; H.S., and R.M-P. reviewed and edited.

