## Supplementary material for "Predictability in the evolution of Orthopteran cardenolide insensitivity": Table S1, Table S2, Table S3, Table S4, Figure S1, Figure S2, Figure S3, Figure S4.

**Supplementary materials**

### Table of Contents

#### 1. Supplementary Tables

**Table S1.** Summary of sites in ATPα1 implicated in cardenolide-binding by site-directed mutagenesis and structural studies.

**Table S2.** List of species and collection information.

**Table S3.** Raw RNA-seq read counts of ATPα1A/B of *Poekilocerus pictus*.

**Table S4.** Probability of positively selected sites under different models.

#### 2. Supplementary Figures

**Fig. S1.** Amino acid sites implicated in cardenolide-binding on ATP1A1 crystal structure.

**Fig. S2.** Amino acid substitutions at the 41 sites implicated in cardenolide-binding.

**Fig. S3.** Relative expression of ATPα1A-D in the head and gut of large milkweed bugs (*Oncopeltus fasciatus*).

**Fig. S4.** Distributions of lineage-specific omega (dN/dS) estimates for ATPα1 excluding the 41 implicated in cardenolide-sensitivity.

#### 3. Supplementary References

**Table S1 Summary of sites in ATPα1 implicated in cardenolide-sensitivity by site-directed mutagenesis and structural studies.**

| Amino Acid Position | Location | Ancestral State | Derived States | Fold Change | References |
|---|---|---|---|---|---|
| 104 | H1 | C | F, A, Y | 6.3-500 | [1,2] |
| 108 | H1 | Y | A, D | 9.1 | [2] |
| 111 | H1-H2 | Q | R, H, L, V, T | 3-12.5 | [3–5] |
| 112 | H1-H2 | A | T, V, K | - | [6] |
| 114 | H1-H2 | T | M, S | - | [6,7] |
| 115 | H1-H2 | S | D | - | [6,8] |
| 116 | H1-H2 | E | D, L | - | [6,8] |
| 117 | H1-H2 | E | * | - | [8] |
| 118 | H1-H2 | P | K, F | 4.4 | [2] |
| 119 | H1-H2 | A | S | - | [7] |
| 120 | H1-H2 | D | R | - | [9] |
| 121 | H1-H2 | D | E, N, A, S | 25-60 | [10,11] |
| 122 | H1-H2 | N | H, D | 8-25 | [3–5,12] |
| 308 | H3-H4 | Y | C, F | 1.5-2.4 | [13] |
| 312 | H3-H4 | D | R | - | [14] |
| 314 | H4 | V | M | - | [14] |
| 315 | H4 | I | V | - | [14] |
| 316 | H4 | F | * | - | [8] |
| 319 | H4 | G | A, S | - | [14,15] |
| 322 | H4 | V | * | - | [8] |
| 323 | H4 | A | * | - | [8] |
| 324 | H4 | N | Y | - | [14] |
| 330 | H4 | L | Q | 7.9 | [16] |
| 331 | H4 | A | G | 3.1 | [16] |
| 338 | H4 | T | N, A | 3.2-3.3 | [16] |
| 367 | H4-H5 | C | * | - | [17] |
| 656 | H4-H5 | A | * | - | [17] |
| 778 | H5 | P | A | - | [15,18] |
| 783 | H5 | F | Y | - | [14] |
| 786 | H5 | F | N, I | 11-19 | [16,19] |
| 793 | H5-H6 | L | P, N, K | 2.8-118 | [12, 20] |
| 797 | H5-H6 | T | A, V, N, C, S | 40-67 | [13,19,21,22] |
| 802 | H6 | C | F | - | [16,20] |
| 804 | H6 | D | E | - | [22] |
| 863 | H7 | F | L | 5.7 | [12] |
| 874 | H7-H8 | K | S | - | [7] |
| 880 | H7-H8 | R | P | 7.9 | [23] |
| 886 | H7-H8 | K | * | - | [8] |
| 898 | H7-H8 | E | E | - | [7] |
| 972 | H9-H10 | R | * | - | [8] |
| 982 | H10 | F | S | 6.3 | [17] |

* Revealed by structural prediction.

**Table S2. List of species and collection information**

| Species | Sex | Life Stage | Tissue | Location | Collected by |
|---|---|---|---|---|---|
| *Aularches miliaris* | F | Adult | Head, muscle, gut, 3 biological replicates | Honey Valley, Karnataka, India | N. Achari, R. Deshmukh |
| *Taphronota calliparea* | F | Adult | Gut, antennae, mouthparts | Sofala, Cheringoma Plateu, Mozambique | R. Mariño-Pérez |
| *Dictyophorus griseus* | F | Adult | Gut, antennae, mouthparts | Sofala, Cheringoma Plateu, Mozambique | R. Mariño-Pérez |
| *Chrotogonus hemipterus* | F | Adult | Head | Sofala, Gorongosa, Mozambique | R. Mariño-Pérez, B. Foquet |
| *Atractomorpha acutipennis* | F | Adult | Head | Sofala, Gorongosa, Mozambique | R. Mariño-Pérez, B. Foquet |
| *Zonocerus elegans* | M | Adult | Head | KwaZulu-Natal, Mpophomeni Hill, South Africa | H. Song, G. Cowper, R. Mariño-Pérez, A. Gomez |
| *Poekilocerus pictus* | M + F | Adult | Head, muscle, gut, 6 biological replicates | Chitoor, Andhra Pradesh, India | N. Achari, R. Deshmukh |
| *Phymateus leprosus* | unknown | Nymph | Head, muscle, gut | KwaZulu-Natal, Ntunjambili Forest, South Africa | H. Song, G. Cowper, R. Mariño-Pérez, A. Armstrong, A. Gomez |
| *Sphenarium purpurascens* | F | Adult | Head | Oaxaca, Near La Colorada, Mexico | R. Mariño-Pérez, B. Foquet, S. Sannabria-Urban, M.E. Pocco |
| *Ochrophlebia cafra* | F | Adult | Head | KwaZulu-Natal, Hlabisa, South Africa | H. Song, G. Cowper, R. Mariño-Pérez, A. Armstrong, A. Gomez |

**Table S3. Raw RNA-seq read counts of ATPα1A/B of *Poekilocerus pictus***

| | Individual | #1 | #2 | #3 | #4 | #5 | #6 |
|---|---|---|---|---|---|---|---|
| **Head** | **ATPα1A** | 1870 | 2145 | 1573 | 455 | 227 | 1365 |
| | **ATPα1B** | 52 | 1397 | 1283 | 14 | 43 | 95 |
| | **# mapped** | 7933248 | 11522005 | 15682417 | 18295987 | 7587412 | 14911717 |
| **Foregut** | **ATPα1A** | 888 | 2943 | 457 | 975 | 3142 | 1022 |
| | **ATPα1B** | 21 | 70 | 17 | 19 | 46 | 18 |
| | **# mapped reads** | 17148883 | 12282284 | 5196066 | 5702546 | 11186498 | 6746261 |
| **Thorax** | **ATPα1A** | 1641 | 1973 | 706 | 588 | 714 | 252 |
| | **ATPα1B** | 46 | 344 | 177 | 298 | 22 | 40 |
| | **# mapped reads** | 11435872 | 12950380 | 11050566 | 8775458 | 24748961 | 20277512 |

**Table S4. Probability of positively selected sites under different models**

| Model No. | Model description | Schematic of model | Positively selected sites | BEB Prob. |
|---|---|---|---|---|
| 1 | Selection in all Apocynaceae-specialist lineages; | 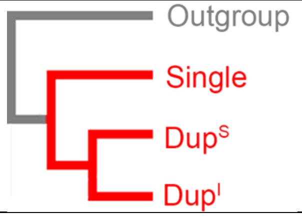  | **111 Q** | **1.000*** |
| 2 | Selection on all Apocynaceae-specialist lineages with single copies of ATPα1 (including ancestral lineages prior to duplication); | 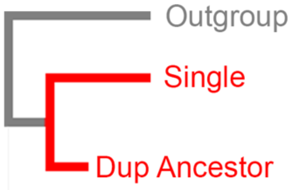  | **111 Q** | **1.000*** |
| 3 | Selection on all Apocynaceae-specialist lineages with duplicate copies of ATPα1; | 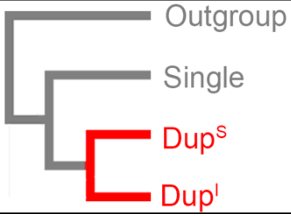  | 301V<br>566 C | 0.966*<br>0.952* |
| 4 | Selection on all putatively less-insensitive ATPα1 duplicate lineages | 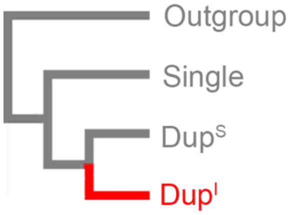  | 102 I<br>106 I<br>**115 V**<br>**118 P**<br>**122 N**<br>163 T<br>177 D<br>274 E<br>**315 I**<br>428 P<br>431 D<br>460 V<br>560 V<br>563 K<br>566 C<br>667 N<br>787 I<br>790 D | 0.953*<br>0.664<br>**0.956***<br>**0.998***<br>**0.993***<br>0.997<br>0.787<br>0.568<br>**0.852**<br>0.553<br>0.530<br>0.690<br>0.980*<br>0.991*<br>0.996*<br>0.981*<br>0.659<br>0.544 |
| 5 | Selection on all outgroup lineages. | 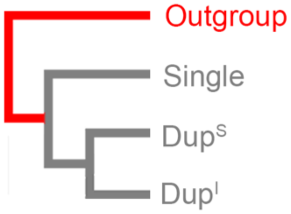  | None | None |

NOTE – Asterisks denote BEB posterior probability >0.95 Sites implicated in cardenolide-sensitivity are highlighted in bold. See Methods for more detail on models.

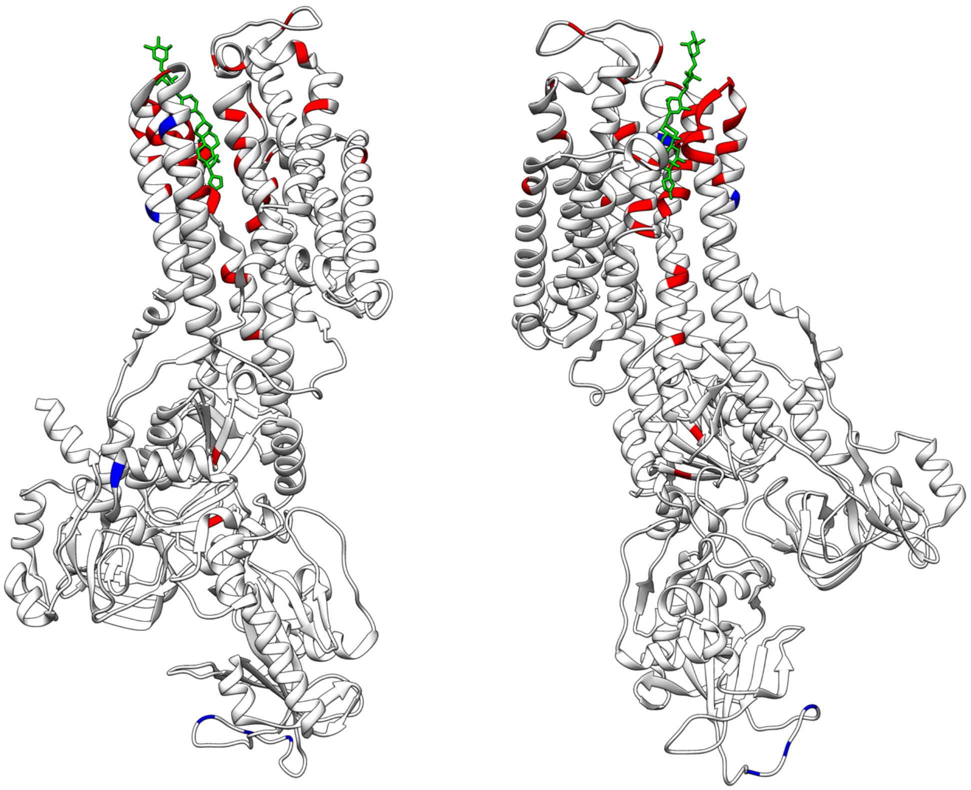

**Figure S1.** Amino acid sites implicated in cardenolide-sensitivity shown on the ATP1A1 crystal structure. 41 amino acid sites (red) implicated in cardenolide-sensitivity are scattered throughout ATP1A1 (*Sus scrofa*, PDB 4RET). Sites in blue are additional sites that have been identified as positively-selected under one of the selection models evaluated in this study (**Figure 7** and **Table S5**). The bound cardenolide digoxin is shown in green.

**Figure S2. Amino acid substitutions at 41 sites of ATPα1 implicated in cardenolide-sensitivity.**

| positions | 104 | 108 | 111 | 112 | 114 | 115 | 116 | 117 | 118 | 119 | 120 | 121 | 122 | 308 | 312 | 314 | 315 | 316 | 319 | 322 | 323 | 324 | 330 | 331 | 338 | 367 | 656 | 778 | 783 | 786 | 787 | 793 | 797 | 802 | 804 | 863 | 874 | 880 | 886 | 898 | 972 | 982 |
|---|---|---|---|---|---|---|---|---|---|---|---|---|---|---|---|---|---|---|---|---|---|---|---|---|---|---|---|---|---|---|---|---|---|---|---|---|---|---|---|---|---|---|
| **Reference** | **C** | **Y** | **Q** | **A** | **T** | V | E | E | **P** | **A** | **D** | **D** | **N** | **Y** | **D** | **V** | **I** | F | **G** | V | **A** | **N** | **L** | **A** | **T** | C | A | P | **F** | **F** | **I** | **L** | **T** | C | **D** | **F** | **L** | **R** | K | E | R | **L** |
| *Mantis religiosa* | . | . | . | . | . | . | . | . | . | . | . | . | . | . | . | . | . | . | . | . | . | . | . | . | . | . | . | . | . | . | . | . | . | . | . | . | . | . | . | . | . | . |
| **Orthopera** | | | | | | | | | | | | | | | | | | | | | | | | | | | | | | | | | | | | | | | | | | |
| *Gryllus firmus* | . | . | . | . | . | . | . | D | . | S | . | . | . | . | . | . | . | . | . | . | . | . | . | . | . | . | . | . | . | . | . | . | . | . | . | . | . | . | . | . | . | . |
| *Tetrix japonica* | . | . | . | . | . | . | . | . | . | . | . | . | . | . | . | . | . | . | . | . | . | . | . | . | . | . | . | . | . | . | . | . | . | . | . | . | . | . | . | . | . | . |
| *Romalea microptera* | . | . | L | . | . | . | . | . | . | S | . | . | . | . | . | . | . | . | . | . | . | . | . | . | . | . | . | . | . | . | . | . | . | . | . | . | . | . | . | . | . | . |
| *Oedaleus asiaticus* | . | . | L | . | . | . | . | . | . | S | . | . | . | . | . | . | . | . | . | . | . | . | . | . | . | . | . | . | . | . | . | . | . | . | . | . | . | . | . | . | . | . |
| *Locusta migratoria* | . | . | L | . | . | . | . | . | . | S | . | . | . | . | . | . | . | . | . | . | . | . | . | . | . | . | . | . | . | . | . | . | . | . | . | . | . | . | . | . | . | . |
| *Aularches miliaris* | . | . | L | . | . | . | . | . | . | S | . | . | . | . | . | . | . | . | . | . | . | . | . | . | . | . | . | . | . | . | . | . | . | . | . | . | . | . | . | . | . | . |
| *Taphronota calliparea* | . | . | L | . | . | . | . | . | . | S | . | . | . | . | . | . | . | . | . | . | . | . | . | . | . | . | . | . | . | . | . | . | . | . | . | . | M | . | . | . | . | . |
| *Dictyophorus griseus* | . | . | L | . | . | . | . | . | . | S | . | . | . | . | . | . | . | . | . | . | . | . | . | . | . | . | . | . | . | . | . | . | . | . | . | . | . | . | . | . | . | . |
| *Chrotogonus hemipterus* | . | . | L | . | . | . | . | . | . | S | . | . | . | . | . | . | . | . | . | . | . | . | . | . | . | . | . | . | . | . | . | . | . | . | . | . | . | . | . | . | . | . |
| *Atractomorpha acutipenni* | . | . | L | . | . | . | . | . | . | S | . | . | . | . | . | . | . | . | . | . | . | . | . | . | . | . | . | . | . | . | . | . | . | . | . | . | . | . | . | . | . | . |
| *Ochrophlebia cafra* | . | . | L | . | . | . | . | . | . | S | . | . | . | . | . | . | . | . | . | . | . | . | . | . | . | . | . | . | . | . | . | . | . | . | . | . | M | . | . | . | . | . |
| *Sphenarium purpurascens* | . | . | L | . | . | . | . | . | . | S | . | . | . | . | . | . | . | . | . | . | . | . | . | . | . | . | . | . | . | . | . | . | . | . | . | . | . | . | . | . | . | . |
| *Zonocerus elegans* | . | . | L | . | . | . | . | . | . | S | . | . | . | . | . | . | . | . | . | . | . | . | . | . | . | . | . | . | . | . | . | . | . | . | . | . | . | . | . | . | . | . |
| *Phymateus leprosus B* | . | . | L | . | . | . | . | . | . | S | . | . | . | . | . | . | . | . | . | . | . | . | . | . | . | . | . | . | . | . | . | . | . | . | . | . | . | . | . | . | . | . |
| *Poekilocerus pictus B* | . | . | L | . | . | . | . | . | . | S | . | . | . | . | . | . | . | . | . | . | . | . | . | . | . | . | . | . | . | . | . | . | . | . | . | . | . | . | . | . | . | . |
| *Phymateus leprosus A* | . | . | L | . | . | T | . | . | S | T | E | N | H | . | . | . | V | . | . | . | . | . | . | . | . | . | . | . | . | . | M | . | . | . | . | . | M | . | . | . | . | . |
| *Poekilocerus pictus A* | . | . | L | . | . | T | . | . | S | S | . | . | H | . | . | . | V | . | . | . | . | . | . | . | . | . | . | . | . | . | M | . | . | . | . | . | . | . | . | . | . | . |
| **Hemiptera** | | | | | | | | | | | | | | | | | | | | | | | | | | | | | | | | | | | | | | | | | | |
| *Oncopeltus fasciatus A* | . | . | T | S | . | . | . | . | A | S | . | N | H | . | . | . | V | . | . | . | . | . | . | . | . | . | . | . | . | N | V | . | A | . | . | . | E | . | . | . | Q | . |
| *Lygaeus kalmii A* | . | . | T | S | . | . | . | . | A | S | . | N | H | . | . | . | V | . | . | . | . | . | . | . | . | . | . | . | . | N | V | . | A | . | . | . | M | . | . | . | Q | . |
| *Oncopeltus fasciatus B* | . | . | T | S | . | E | . | . | S | S | . | . | H | . | . | . | V | . | . | . | . | . | . | . | . | . | . | . | . | N | V | . | . | . | . | . | M | . | . | . | . | . |
| *Lygaeus kalmii B* | . | . | T | S | . | E | . | . | S | S | . | . | H | . | . | . | V | . | . | . | . | . | . | . | . | . | . | . | . | N | V | . | . | . | . | . | M | . | . | . | . | . |
| *Oncopeltus fasciatus C* | . | . | . | . | . | . | . | . | . | S | . | . | H | . | . | . | . | . | . | . | . | . | . | . | . | . | . | . | . | . | M | . | S | . | . | . | M | . | . | . | . | . |
| *Lygaeus kalmii C* | . | . | . | . | . | . | . | . | . | S | . | . | H | . | . | . | . | . | . | . | . | . | . | . | . | . | . | . | . | . | M | . | S | . | . | . | M | . | . | . | . | . |
| *Oncopeltus fasciatus D* | . | . | . | . | . | . | . | . | . | S | . | . | H | . | . | . | . | . | . | . | . | . | . | . | . | . | . | . | . | . | . | . | . | . | . | . | V | . | . | . | . | . |
| *Boisea trivittata* | . | . | . | . | . | . | . | . | . | S | . | . | . | . | . | . | . | . | . | . | . | . | . | . | . | . | . | . | . | . | . | . | . | . | . | . | T | . | . | . | . | . |
| *Pyrrhocoris apterus* | . | . | . | . | . | . | . | . | . | S | . | . | . | . | . | . | . | . | . | . | . | . | . | . | . | . | . | . | . | . | . | . | . | . | . | - | - | - | - | - | - | - |
| *Largus sp.* | . | . | E | T | . | . | . | . | . | S | . | . | . | . | . | . | . | . | . | . | . | . | . | . | . | . | . | . | . | . | . | . | . | . | . | - | - | - | - | - | - | - |
| *Bemisia tabaci* | . | . | E | S | . | I | . | . | . | . | . | . | . | . | E | . | . | . | . | . | . | . | . | . | . | . | . | . | . | . | V | . | V | . | . | . | M | . | . | . | . | . |
| *Acyrthosiphon pisum* | . | H | E | T | . | T | . | D | . | N | . | . | Y | C | . | I | . | . | . | . | . | A | . | . | . | . | . | . | . | . | . | . | A | . | . | . | Y | . | R | . | . | I |
| *Aphis nerii* | . | H | E | T | . | T | . | D | . | N | . | . | Y | C | . | I | . | . | . | . | . | A | . | . | . | . | . | . | . | . | . | . | A | . | . | . | Y | . | R | . | . | I |
| **Hymenoptera** | | | | | | | | | | | | | | | | | | | | | | | | | | | | | | | | | | | | | | | | | | |
| *Monophadnus latus* | . | . | T | . | . | L | . | . | S | N | . | . | H | . | . | . | . | . | . | . | . | . | . | . | . | . | . | . | . | . | M | . | . | . | . | - | - | - | - | - | - | - |
| *Pachyprotasis variegata* | . | . | . | . | . | . | . | D | . | N | . | . | . | . | . | . | . | . | . | . | . | . | . | . | . | . | . | . | . | . | . | . | . | . | . | - | - | - | - | - | - | - |
| *Apis mellifera* | . | . | . | . | . | S | . | D | . | N | . | . | . | . | . | . | . | . | . | . | . | . | . | . | . | . | . | . | . | . | . | . | . | . | . | - | - | - | - | - | - | - |
| **Coleoptera** | | | | | | | | | | | | | | | | | | | | | | | | | | | | | | | | | | | | | | | | | | |
| *Chrysochus auratus A* | . | . | L | . | . | . | . | . | . | S | . | . | . | . | . | . | . | . | . | . | . | . | . | . | . | . | . | . | . | . | . | . | . | . | . | . | M | . | . | . | . | . |
| *Chrysochus auratus B* | . | . | V | V | . | . | . | . | A | S | . | . | H | . | . | . | L | . | . | . | . | . | . | . | . | . | . | . | . | . | M | . | . | . | . | . | M | S | . | . | . | . |
| *Chrysochus cobalitinus A* | . | . | L | . | . | . | . | . | . | S | . | . | . | . | . | . | . | . | . | . | . | . | . | . | . | . | . | . | . | . | . | . | . | . | . | . | M | . | . | . | . | . |
| *Chrysochus cobalitinus B* | . | . | V | V | . | . | . | . | A | S | . | . | H | . | . | . | L | . | . | . | . | . | . | . | . | . | . | . | . | . | M | . | . | . | . | . | M | . | . | . | . | . |
| *Chrysochus asclepiadeus* | . | . | L | . | . | . | . | . | . | S | . | . | . | . | . | . | . | . | . | . | . | . | . | . | . | . | . | . | . | . | M | . | . | . | . | - | - | - | - | - | - | - |
| *Labidomera clivicolis* | . | . | V | . | . | A | . | . | . | N | . | . | H | . | . | . | . | . | . | . | . | . | . | . | . | . | . | . | . | . | . | . | . | . | . | . | M | . | . | . | . | . |
| *Plagiodera versicolora* | . | . | . | . | . | . | . | . | . | . | . | . | . | . | . | . | . | . | . | . | . | . | . | . | . | . | . | . | . | . | . | . | . | . | . | . | M | . | . | . | . | . |
| *Gastrophysa viridula* | . | . | . | . | . | . | . | . | . | . | . | . | . | . | . | . | . | . | . | . | . | . | . | . | . | . | . | . | . | . | . | . | . | . | . | . | M | . | . | . | - | - |
| *Alticini sp.* | . | . | . | . | . | . | . | . | . | . | . | . | . | . | . | . | . | . | . | . | . | . | . | . | . | . | . | . | . | . | . | . | . | . | . | - | - | - | - | - | - | - |
| *Tetraopes tetraophthalmu* | . | . | L | . | . | . | . | . | . | S | . | . | . | . | . | . | V | . | . | . | . | . | . | . | . | . | . | . | . | . | . | . | . | . | . | . | R | . | . | . | . | . |
| *Megacyllene robiniae* | . | . | . | . | . | . | . | . | . | . | . | . | . | . | . | . | . | . | . | . | . | . | . | . | . | . | . | . | . | . | . | . | . | . | . | . | R | . | . | . | . | . |
| *Rhyssomatus lineaticollis B* | Y | . | . | . | . | . | . | . | . | S | . | . | Y | . | . | . | . | . | . | . | . | . | . | . | . | . | . | . | . | . | . | . | . | . | . | . | M | . | . | . | . | . |
| *Rhyssomatus lineaticollis A* | . | . | T | . | . | . | . | . | . | S | . | . | . | . | . | . | . | . | . | . | . | . | . | . | . | . | . | . | . | . | . | . | . | . | . | . | M | . | . | . | . | . |
| *Cyrtepistomus castaneus* | . | . | . | . | . | . | . | . | . | . | . | . | . | . | . | . | . | . | . | . | . | . | . | . | . | . | . | . | . | . | . | . | . | . | . | . | M | . | . | . | . | . |
| *Tribolium castaneum* | . | . | . | . | . | . | . | . | . | . | . | . | . | . | . | . | . | . | . | . | . | . | . | . | . | . | . | . | . | . | . | . | . | . | . | . | T | . | . | . | . | . |

*(Continues on the next page...)*

(...continued)

| positions | **104** | **108** | **111** | **112** | **114** | 115 | 116 | 117 | **118** | **119** | **120** | **121** | **122** | **308** | **312** | **314** | **315** | 316 | **319** | 322 | **323** | **324** | **330** | **331** | **338** | 367 | 756 | 778 | **783** | **786** | **787** | **793** | **797** | 802 | **804** | **863** | **874** | **880** | 886 | 898 | 972 | **982** |
|---|---|---|---|---|---|---|---|---|---|---|---|---|---|---|---|---|---|---|---|---|---|---|---|---|---|---|---|---|---|---|---|---|---|---|---|---|---|---|---|---|---|---|
| **Reference** | **C** | **Y** | **Q** | **A** | **T** | V | E | E | **P** | **A** | **D** | **D** | **N** | **Y** | **D** | **V** | **I** | F | **G** | V | **A** | **N** | **L** | **A** | **T** | C | A | P | **F** | **F** | **I** | **L** | **T** | C | **D** | **F** | **L** | **R** | K | E | R | **L** |
| **Diptera** | | | | | | | | | | | | | | | | | | | | | | | | | | | | | | | | | | | | | | | | | | |
| *Lucilia bufonivora* | . | . | . | . | . | S | . | . | . | S | . | . | . | . | . | . | . | . | . | . | . | . | . | . | . | . | . | . | . | . | . | . | . | . | . | - | - | - | - | - | - | - |
| *Drosophila melanogaster* | . | . | . | . | . | S | . | . | . | . | . | . | . | . | . | . | . | . | . | . | . | . | . | . | . | . | . | . | . | . | . | . | . | . | . | . | K | . | . | . | . | F |
| *Drosophila guanche* | . | . | . | . | . | S | . | . | . | S | . | . | . | . | . | . | . | . | . | . | . | . | . | . | . | . | . | . | . | . | . | . | . | . | . | . | R | . | . | . | . | F |
| *Drosophila subobscura* | . | . | . | . | . | S | . | . | . | S | . | . | . | . | . | . | . | . | . | . | . | . | . | . | . | . | . | . | . | . | . | . | . | . | . | . | R | . | . | . | . | T |
| *Liriomyza eupatorii* | . | . | . | . | . | S | . | . | . | S | . | . | . | . | . | . | . | . | . | . | . | . | . | . | . | . | . | . | . | . | . | . | . | . | . | - | - | - | - | - | - | - |
| *Liriomyza asclepiadis* | . | . | . | . | . | S | . | . | . | S | . | . | H | . | . | . | . | . | . | . | . | . | . | . | . | . | . | . | . | . | . | . | . | . | . | . | R | . | . | . | . | . |
| *Liriomyza sp.* | . | . | L | . | . | T | . | . | . | S | . | . | . | . | . | . | . | . | . | . | . | . | . | . | . | . | . | . | . | . | . | . | . | . | . | - | - | - | - | - | - | - |
| *Chromatomyia horticola* | . | . | . | . | . | S | . | . | . | S | . | . | . | . | . | . | . | . | . | . | . | . | . | . | . | . | . | . | . | . | . | . | . | . | . | - | - | - | - | - | - | - |
| *Phytomyza fallaciosa* | . | . | . | . | . | S | . | . | . | S | . | . | . | . | . | . | . | . | . | . | . | . | . | . | . | . | . | . | . | . | . | . | . | . | . | - | - | - | - | - | - | - |
| *Phytomyza hellebori A* | . | . | H | . | . | S | . | . | . | S | . | . | H | . | . | . | . | . | . | . | . | . | . | . | . | . | . | . | . | . | M | . | . | . | . | - | - | - | - | - | - | - |
| *Phytomyza hellebori B* | . | . | . | . | . | S | . | . | . | S | . | . | . | . | . | . | . | . | . | . | . | . | . | . | . | . | . | . | . | . | M | . | . | . | . | - | - | - | - | - | - | - |
| *Phytomyza digitalis* | . | . | L | . | . | S | . | . | . | S | . | . | . | . | . | . | . | . | . | . | . | . | . | . | . | . | . | . | . | . | . | . | . | . | . | - | - | - | - | - | - | - |
| *Phytomyza crassiseta* | . | . | . | . | . | S | . | . | . | S | . | . | . | . | . | . | . | . | . | . | . | . | . | . | . | . | . | . | . | . | . | . | . | . | . | - | - | - | - | - | - | - |
| *Phytomyza ilicis* | . | . | . | . | . | S | . | . | . | S | . | . | . | . | . | . | . | . | . | . | . | . | . | . | . | . | . | . | . | . | . | . | . | . | . | - | - | - | - | - | - | - |
| *Napomyza scrophulariae* | . | . | L | . | . | E | . | . | . | S | . | . | . | . | . | . | . | . | . | . | . | . | . | . | . | . | . | . | . | . | . | . | . | . | . | - | - | - | - | - | - | - |
| *Napomyza lateralis* | . | . | L | . | . | E | . | . | . | S | . | . | . | . | . | . | . | . | . | . | . | . | . | . | . | . | . | . | . | . | . | . | . | . | . | - | - | - | - | - | - | - |
| **Lepidoptera** | | | | | | | | | | | | | | | | | | | | | | | | | | | | | | | | | | | | | | | | | | |
| *Danaus plexippus* | . | . | V | . | . | . | . | . | . | S | . | . | H | . | . | . | . | . | . | . | . | . | . | . | . | . | . | . | . | . | . | . | . | . | . | . | T | . | . | . | . | . |
| *Danaus erippus* | . | . | V | . | . | . | . | . | . | S | . | . | H | . | . | . | . | . | . | . | . | . | . | . | . | . | . | . | . | . | . | . | . | . | . | - | - | - | - | - | - | - |
| *Danaus eresimus* | . | . | V | . | . | . | . | . | . | S | . | . | H | . | . | . | . | . | . | . | . | . | . | . | . | . | . | . | . | . | . | . | . | . | . | . | T | . | . | . | . | . |
| *Danaus chrysippus* | . | . | V | . | . | . | . | . | . | S | . | . | . | . | . | . | . | . | . | . | . | . | . | . | . | . | . | . | . | . | . | . | . | . | . | . | T | . | . | . | . | . |
| *Danaus gilippus* | . | . | V | . | . | . | . | . | . | S | . | . | . | . | . | . | . | . | . | . | . | . | . | . | . | . | . | . | . | . | . | . | . | . | . | . | T | . | . | . | . | . |
| *Danaus genutia* | . | . | V | . | . | . | . | . | . | S | . | . | . | . | . | . | . | . | . | . | . | . | . | . | . | . | . | . | . | . | . | . | . | . | . | . | T | . | - | - | - | - |
| *Tirumala limniace* | . | . | V | . | . | . | . | . | . | S | . | . | . | . | . | . | . | . | . | . | . | . | . | . | . | . | . | . | . | . | . | . | . | . | . | . | V | - | - | - | - | - |
| *Tirumala petiverana* | . | . | V | . | . | . | . | . | . | S | . | . | . | . | . | . | . | . | . | . | . | . | . | . | . | . | . | . | . | . | . | . | . | . | . | . | V | . | - | - | - | - |
| *Tirumala septentrionis* | . | . | V | . | . | . | . | . | . | S | . | . | . | . | . | . | . | . | . | . | . | . | . | . | . | . | . | . | . | . | . | . | . | . | . | . | V | . | - | - | - | - |
| *Euploea core core* | . | . | L | . | . | . | . | . | . | S | . | . | . | . | . | . | . | . | . | . | . | . | . | . | . | . | . | . | . | . | . | . | . | . | . | . | M | . | . | . | . | . |
| *Euploea phaenareta* | . | . | L | . | . | . | . | . | . | S | . | . | . | . | . | . | . | . | . | . | . | . | . | . | . | . | . | . | . | . | . | . | . | . | . | . | . | . | - | - | - | - |
| *Idea leuconoe* | . | . | L | . | . | . | . | . | . | S | . | . | . | . | . | . | . | . | . | . | . | . | . | . | . | . | . | . | . | . | . | . | . | . | . | . | M | . | - | - | - | - |
| *Parantica aglea* | . | . | L | . | . | . | . | . | . | S | . | . | . | . | . | . | . | . | . | . | . | . | . | . | . | . | . | . | . | . | . | . | . | . | . | - | - | - | - | - | - | - |
| *Ideopsis juventa* | . | . | L | . | . | . | . | . | . | S | . | . | . | . | . | . | . | . | . | . | . | . | . | . | . | . | . | . | . | . | . | . | . | . | . | - | - | - | - | - | - | - |
| *Mechanitis polymnia* | . | . | L | . | . | . | . | . | . | . | . | . | . | . | . | . | . | . | . | . | . | . | . | . | . | . | . | . | . | . | . | . | . | . | . | . | T | . | - | - | - | - |
| *Lycorea halia* | . | . | L | . | . | . | . | . | . | S | . | . | . | . | . | . | . | . | . | . | . | . | . | . | . | . | . | . | . | . | . | . | . | . | . | . | T | . | . | . | . | . |
| *Limenitis archippus* | . | . | . | . | . | . | . | . | . | . | . | . | . | . | . | . | . | . | . | . | . | . | . | . | . | . | . | . | . | . | . | . | . | . | . | . | . | . | . | . | . | . |
| *Papilio glaucus* | . | . | . | . | . | . | . | . | . | . | . | . | . | . | . | . | . | . | . | . | . | . | . | . | . | . | . | . | . | . | . | . | . | . | . | . | . | . | . | . | . | . |
| *Cycnia oregonensis* | . | . | . | . | . | . | . | . | . | . | . | . | . | . | . | . | . | . | . | . | . | . | . | . | . | . | . | . | . | . | . | . | . | . | . | - | - | - | - | - | - | - |
| *Cycnia tenera* | . | . | L | . | . | . | . | . | . | S | . | . | . | . | . | . | . | . | . | . | . | . | . | . | . | . | . | . | . | . | . | . | . | . | . | . | M | . | . | . | . | . |
| *Lerina incarnata* | . | . | L | . | . | . | . | . | . | S | . | . | . | . | . | . | L | . | . | . | . | . | . | . | . | . | . | . | . | . | . | . | . | - | - | - | - | - | - | - | - | - |
| *Pygoctenucha terminalis* | . | . | L | . | . | . | . | . | . | S | . | . | . | . | . | . | . | . | . | . | . | . | . | . | . | . | . | . | . | . | . | . | . | . | . | - | - | - | - | - | - | - |
| *Lophocampa caryae* | . | . | L | . | . | . | . | . | . | . | . | . | . | . | . | . | . | . | . | . | . | . | . | . | . | . | . | . | . | . | . | . | . | . | . | . | . | . | . | . | . | . |
| *Empyreuma pugione* | . | . | L | . | . | . | . | . | . | . | . | . | . | . | . | . | . | . | . | . | . | . | . | . | . | . | . | . | . | . | . | . | . | . | . | - | - | - | - | - | - | - |
| *Olepa ricini* | . | . | . | . | . | . | . | . | . | . | . | . | . | . | . | . | . | . | . | . | . | . | . | . | . | . | . | . | . | . | . | . | . | . | . | . | . | . | . | . | . | . |
| *Trichordestra legitima* | . | . | . | . | . | . | . | . | . | . | . | . | . | . | . | . | . | . | . | . | . | . | . | . | . | . | . | . | . | . | . | . | . | . | . | . | M | . | . | . | . | . |
| *Saucrobotys futilalis* | . | . | L | . | . | . | . | . | . | S | . | . | . | . | . | . | . | . | . | . | . | . | . | . | . | . | . | . | . | . | . | . | . | . | - | - | - | - | - | - | - | - |
| *Daphnis nerii* | . | . | L | . | . | . | . | . | . | S | . | . | . | . | . | . | . | . | . | . | . | . | . | . | . | . | . | . | . | . | . | . | . | . | . | . | M | . | . | . | . | . |
| *Bombyx mori* | . | . | . | . | . | . | . | . | . | . | . | . | . | . | . | . | . | . | . | . | . | . | . | . | . | . | . | . | . | . | . | . | . | . | . | . | M | . | . | . | . | . |

NOTE: Reference is the consensus sequence of non-Apocynaceae-specialists. Dots indicate identical to the reference while letters refer to nonsynonymous amino acid changes. Hyphen - indicates missing data.

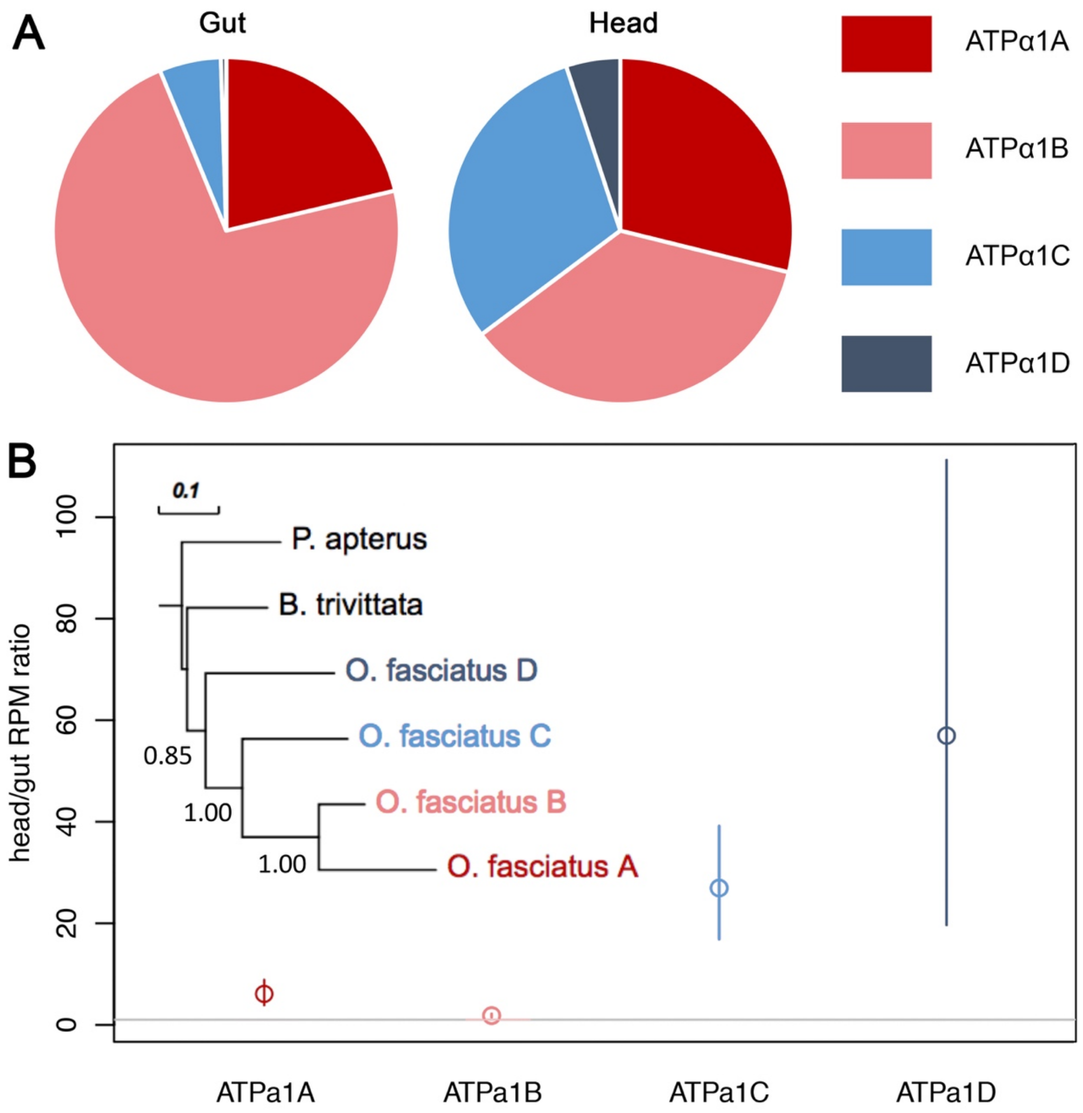


**Figure S3.** Relative expression of four ATPα1 copies in the head and gut of large milkweed bugs (*Oncopeltus fasciatus*). **A)** The average percentages of ATPα1A-D of the total ATPα1 in both gut and head. **B**) The normalised counts (reads per million, or RPM) ratio of head to gut. The grey horizontal line represents the head/gut ratio of 1. Open circles and whiskers represent means and approximate 95% confidence interval determined by bootstrap resampling the count data by individual. The phylogeny was constructed using a maximum likelihood method implemented in SeaView based on nucleotide-coding nucleotide sequences of ATPα1 with bootstrap values shown.

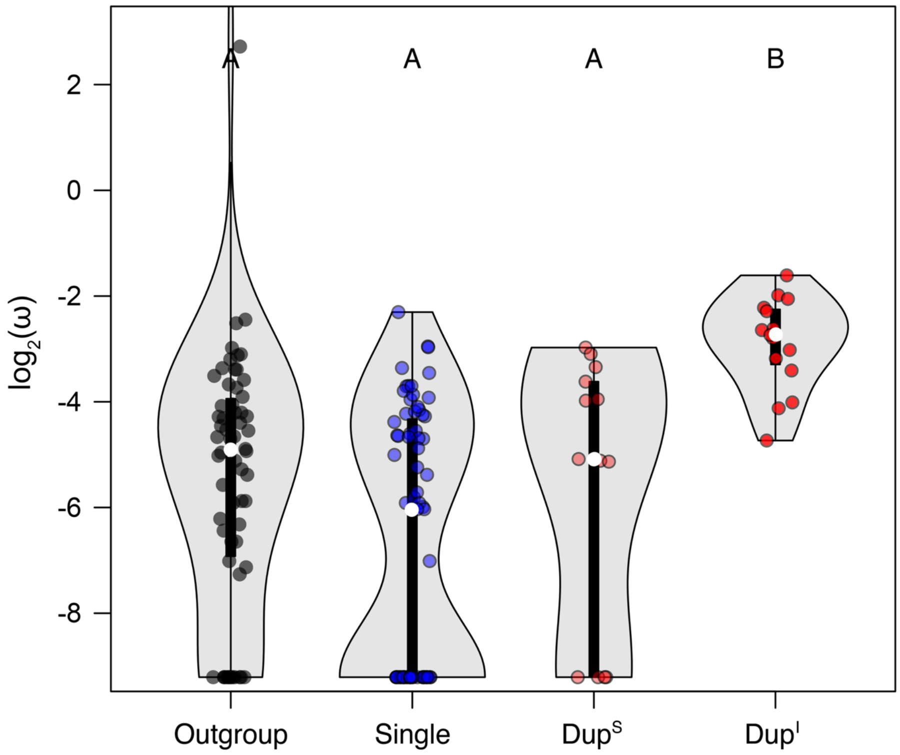


**Figure S4**. Distributions of lineage-specific omega (dN/dS) estimates for ATPα1 excluding the 41 implicated in cardenolide-sensitivity. The distributions of omega estimates are shown with median and 95% confidence intervals. Letters A and B indicate significantly different categories. There is a significant difference between the omega ratios of $Dup^I$ and those of the three other groups ($Dup^I$ vs Outgroup P=6e-5, $Dup^I$ vs Single P=4e-8, $Dup^I$ vs $Dup^S$ P=3e-3). P-values were estimated using Dunn's test of multiple comparisons using rank sums as implement in R (dunn.test) and corrected using the Benjamini-Hochberg method.
